# Detecting Binocular Disparity in Photorealistic Objects

**DOI:** 10.64898/2026.11.27.769657

**Authors:** Pasuparthi Gouri Pravalitha, Andrew J. Parker

## Abstract

Human stereoacuity has been characterised largely with stimuli designed to be monocularly uninformative, such as random dot stereograms, whereas studies using naturalistic images have mostly measured the appearance of clearly visible depth rather than the threshold at which a binocular signal can be detected. We measured binocular disparity detection thresholds for photorealistic objects rendered from high-resolution laser scans, using a spatial four-alternative forced-choice odd-one-out task, in which observers reported which of four objects carried added binocular depth. Depth was parameterised relative to each observer’s interpupillary distance, an observer-relative scaling that can be applied identically across species. Sixteen observers completed a static condition and thirteen also completed a condition in which the objects rotated in depth, providing structure-from-motion in addition to disparity. Detection thresholds were reliably measurable with photorealistic objects, corresponding to a maximum within-object disparity of 0.89 arcmin (0.015° of visual angle) for static objects. Both accuracy and correct-trial reaction times varied systematically with disparity magnitude, while incorrect-trial reaction times did not. In a monocular control, in which the target object was shown with its left-eye image in both eyes, performance fell to chance, confirming that the target was identified by its binocular disparity. Thresholds were substantially higher (1.48 arcmin, 0.025°) and responses slower for rotating compared with static objects, indicating that object motion imposed a measurable cost on the extraction of binocular depth. These results establish a behavioural benchmark and a transferable task design for neurophysiological studies of binocular depth processing in mid-level visual cortex.

## 1. Introduction

The perception of a stable, three-dimensional world is one of the central achievements of the visual system. To recover the depth structure of a scene, the brain draws on multiple, partially redundant, sources of information: binocular disparity, motion parallax, shading, texture gradients, occlusion, and the familiar shapes and sizes of known objects (Howard & Rogers, 2002). A long tradition of psychophysical and computational work has shown that these cues are not used in isolation but are combined, with the influence of each cue weighted according to its reliability in the prevailing viewing conditions (Landy, Maloney, Johnston, & Young, 1995; Johnston, Cumming, & Parker, 1993; Ernst & Banks, 2002; Hillis, Watt, Landy, & Banks, 2004; Hibbard, Asher, & Hornsey, 2025). Within this framework, robust depth perception emerges because one cue can compensate for another when the latter becomes unreliable, and because the simultaneous availability of several cues improves the precision of the resulting depth estimate.

Binocular disparity refers to the small differences between the two eyes’ images that arise from their horizontal separation, and it is among the most precise of these cues. Under optimal laboratory conditions, human observers can discriminate disparities corresponding to a few seconds of arc, a sensitivity characterised in detail using simplified stimuli such as random dot stereograms, bars, and isolated points of light (Cumming & DeAngelis, 2001; Howard & Rogers, 2002; McKee, Levi, & Bowne, 1990). The random dot stereogram in particular has been the workhorse of stereopsis research because it isolates binocular disparity from all monocular form cues: the three-dimensional structure of the stimulus is invisible to either eye alone and emerges only upon binocular fusion. This approach using careful isolation of the disparity cue has been enormously productive, yielding precise estimates of stereoacuity (McKee et al., 1990; Coutant & Westheimer, 1993) and constraining models of the underlying neural computation.

Yet, this experimental purity creates a gap between the conditions under which stereopsis is typically measured and the conditions under which it normally operates. In natural vision, binocular disparity is almost never the only cue to depth. It is embedded within scenes containing shading, texture, occlusion, and recognisable objects whose familiar shapes carry their own depth information (Hibbard, Asher, & Hornsey, 2025; Hibbard, Hornsey, & Asher, 2023; Hornsey, Hibbard, & Scarfe, 2015). Whether the disparity sensitivity measured with sparse, abstract stimuli generalises to such object-rich, photorealistic contexts is not obvious. Additional monocular structure could in principle facilitate disparity processing by providing matchable features and disambiguating the correspondence problem, or it could compete with, and partially mask, the binocular signal, as can occur when cues specify conflicting depth (Johnston, Cumming, & Landy, 1994). Recent work using stereoscopic photographs of natural scenes indicates that binocular information continues to enhance the reliability and apparent magnitude of perceived depth, even when rich pictorial cues are present (Hibbard et al., 2023; Hornsey et al., 2015). Despite this, the threshold sensitivity to disparity in photorealistic objects has received comparatively little direct measurement.

There is, moreover, good reason to expect that the depth recovered from a smoothly curved, fully shaded object is not a simple readout of the disparity field. Observers systematically misperceive the depth of binocularly defined curved surfaces, and the pattern of error depends on surface geometry and viewing distance (Hornsey, Hibbard, & Scarfe, 2016; Hibbard, Goutcher, O’Kane, & Scarfe, 2012). Reverse correlation analysis further shows that observers do not sample the disparity field uniformly when estimating three-dimensional shape, but weight particular surface regions preferentially (Scarfe & Hibbard, 2013). Photorealistic objects of the kind used here are exactly the class of stimulus for which such effects are expected to matter, which makes it an open empirical question whether orderly detection thresholds can be recovered at all.

The contribution of motion adds a further dimension to this question. When an object rotates, the changing pattern of its retinal projection provides a powerful cue to three-dimensional shape, known as structure-from-motion (Howard & Rogers, 2002). Motion and stereopsis interact strongly in the specification of three-dimensional shape: when both cues are present and congruent, the perceptual distortions that each cue produces in isolation are reduced or abolished, consistent with a weighted combination of the two signals after appropriate scaling for viewing distance (Johnston et al., 1993, 1994; Tittle & Braunstein, 1993; Tittle & Perotti, 1997). Against this, motion-in-depth has also been shown to disrupt rather than assist the recovery of metric structure: disparity-defined objects moving in depth do not elicit three-dimensional shape constancy, indicating a lack of automatic compensation during changes of binocular geometry (Scarfe & Hibbard, 2006). How the addition of object motion affects the detectability of a binocular disparity signal, as opposed to the perceived shape of a clearly visible object, therefore remains an open question, and particularly so for photorealistic stimuli.

Addressing these questions has acquired new practical importance. Photorealistic stimuli generated from high-resolution laser scans of real objects can be rendered from arbitrary viewpoints and manipulated quantitatively while retaining their naturalistic appearance, offering a controlled bridge between abstract laboratory stimuli and real-world scenes (Scarfe & Glennerster, 2019, 2015). Such stimuli are increasingly central to programmes that seek to link human psychophysics with neurophysiological recordings in non-human primates, where the goal is to understand how populations of neurons in mid-level visual areas such as V4 integrate binocular depth with other cues in realistic visual contexts (Parker, 2020; Pasupathy, Popovkina, & Kim, 2020; Smith & Parker, 2021). For such a programme to succeed, it is first necessary to establish that reliable, quantitative measures of disparity sensitivity can be obtained with photorealistic objects, using a task that is suitable for both human and macaque observers.

### 1.1 What distinguishes the present study

The present study occupies an unusual position in this literature, because of four features that have not previously been combined in a single experiment.

#### Threshold measurement with photorealistic objects

Almost everything known quantitatively about human stereoacuity derives from stimuli designed to be monocularly uninformative (Cumming & DeAngelis, 2001; McKee et al., 1990; Howard & Rogers, 2002). Conversely, studies using naturalistic or photorealistic images have almost all measured suprathreshold quantities, namely the apparent magnitude, reliability, or ordinal structure of perceived depth in a clearly visible object or scene (Hibbard et al., 2023; Hornsey et al., 2015, 2016; Scarfe & Hibbard, 2013). An important exception is White and Burge (2025), who measured disparity discrimination thresholds for 1° stereo-image patches sampled from natural scenes with laser-measured ground-truth depth, and showed that natural variation in local depth structure raises thresholds multiplicatively. Their stimuli, however, were small foveal patches discriminated against a disparity pedestal, rather than whole, recognisable objects carrying shading, outline and familiar shape. To our knowledge, no previous study has combined photorealistic, laser-scanned objects with a systematic psychometric estimate of the detection threshold for added binocular disparity. This matters because the two measurement regimes could be dissociable: cues that dominate the suprathreshold percept need not limit near threshold detection, and strong pictorial structure could in principle either aid or mask a small binocular signal.

#### Observer-relative parameterisation in %IPD

Depth magnitude here is specified not in absolute angular units at the outset, but as a proportion of each observer’s own interpupillary distance (%IPD), with the stereo pairs pre-rendered at the individual observer’s measured interpupillary distance. This normalises for individual differences in optical geometry, so that a given nominal level corresponds to the same relationship between viewing geometry and rendered depth in every observer. Because interocular separation is the natural scaling variable across species, the same parameterisation can also be applied without modification to other species, whose interpupillary distance differs substantially from that of humans. The %IPD axis therefore provides a common currency in which human and non-human thresholds can be compared directly, which disparity in arcmin does not. Angular values are recovered from the rendering and display geometry (Section 2.4) so that the results remain comparable with the published stereoacuity literature.

#### A paradigm already validated in the macaque

The spatial four-alternative forced-choice odd-one-out task used here is not merely suitable in principle for animal work; it is the same four-quadrant depth change detection paradigm already run in awake, behaving macaques while neuronal populations were recorded simultaneously from V1 and V4 with chronically implanted arrays (Smith & Parker, 2021). In that work, animals detected which of four random dot stereograms underwent a step change in binocular depth and indicated it by a saccade, with orderly psychometric and chronometric functions. The human data reported here are therefore a directly commensurable baseline, obtained with the identical decision structure, chance level, and dependent measures, differing only in the stimulus class and the response modality.

#### Static versus rotating objects within observer

Finally, testing each observer with both static and smoothly rotating objects allows the effect of a second naturalistic depth cue, structure-from-motion, on threshold disparity detection to be assessed within subject. Prior work on the interaction between stereopsis and motion has concentrated on suprathreshold shape judgements and has yielded a mixed picture, with evidence both for cooperative combination (Johnston et al., 1994; Tittle & Braunstein, 1993) and for a failure of shape constancy when the binocular geometry changes over time (Scarfe & Hibbard, 2006). Measuring the interaction at threshold, in objects that carry the full complement of pictorial cues, addresses a regime that neither line of previous work has covered.

Here, we measured binocular disparity detection thresholds for photorealistic objects using a spatial four-alternative forced-choice odd-one-out task, a method that is robust for naive observers and free of some of the response biases that may affect two-alternative judgements (Jäkel & Wichmann, 2006; Vancleef et al., 2018). On each trial, observers viewed four photorealistic stereo image pairs of everyday objects; three distractor objects carried no added binocular depth and one target object carried a disparity signal, with the observer reporting which of the four was different. We asked three questions: whether reliable disparity detection thresholds can be obtained with photorealistic objects; how disparity magnitude affects detection accuracy and reaction time; and how the addition of object motion affects disparity detectability. Beyond their intrinsic psychophysical interest, the answers establish a behavioural foundation and a task design that can be transferred directly to subsequent neurophysiological recordings in macaque visual cortex.

## 2. Methods

### 2.1 Participants

Observers with stereo deficiency, assessed using the TNO test for stereoscopic vision (Walraven, n.d.), were excluded prior to the main experiment. Sixteen adult observers (5 men, 11 women; aged 23–65 years; normal or corrected-to-normal visual acuity of at least 20/20) completed the static condition, and thirteen of these also completed the motion condition; the remaining three (P14–P16) were tested in the static condition only (see Table 1). One observer was an author of the study; all others were naive to its purpose. Interpupillary distance was measured for each participant and ranged from 58 to 63 mm. All participants gave informed consent prior to testing, and the study was conducted in accordance with the Declaration of Helsinki (World Medical Association, 2025) and approved by the local ethics committee.

**Table 1.** Per-observer thresholds and reaction times across the static and motion conditions.

| **Obs** | **IPD (mm)** | **Static thr, %IPD [95% CI]** | **Static thr, arcmin [95% CI]** | **Motion thr, %IPD [95% CI]** | **Motion thr, arcmin [95% CI]** | **Static RT, ms [95% CI]** | **Motion RT, ms [95% CI]** |
|---|---|---|---|---|---|---|---|
| P1 | 63 | 66.8 [58.2, 75.8] | 1.06 [0.92, 1.20] | n.b. | — | 2112 [2010, 2218] | 1499 [1430, 1571] |
| P2 | 63 | 49.8 [41.7, 57.9] | 0.79 [0.66, 0.91] | 99.9 [89.5, 114.1] | 1.58 [1.41, 1.80] | 1793 [1728, 1860] | 2222 [2121, 2319] |
| P3 | 63 | 34.4 [27.6, 40.7] | 0.54 [0.44, 0.64] | 84.7 [74.7, 96.1] | 1.34 [1.18, 1.52] | 1843 [1773, 1916] | 3181 [2988, 3395] |
| P4 | 63 | 50.4 [40.7, 59.7] | 0.80 [0.64, 0.94] | 75.5 [65.5, 85.8] | 1.19 [1.04, 1.36] | 2635 [2489, 2784] | 2991 [2874, 3116] |
| P5 | 63 | 54.4 [46.3, 62.1] | 0.86 [0.73, 0.98] | 89.6 [79.3, 102.3] | 1.42 [1.25, 1.62] | 1971 [1873, 2071] | 2178 [2077, 2277] |
| P6 | 62† | 55.9 [47.8, 64.2] | 0.87 [0.74, 1.00] | 89.3 [78.7, 100.6] | 1.39 [1.22, 1.56] | 2765 [2619, 2927] | 3783 [3593, 3993] |
| P7 | 62† | 42.1 [34.5, 49.1] | 0.66 [0.54, 0.76] | 85.3 [73.5, 99.7] | 1.33 [1.14, 1.55] | 3046 [2699, 3389] | 3320 [3053, 3602] |
| P8 | 58 | 89.5 [76.5, 111.7] | 1.30 [1.11, 1.63] | 131.0 [107.3, 190.2] | 1.91 [1.56, 2.77] | 2681 [2429, 2940] | 3027 [2748, 3349] |
| P9 | 62† | 74.0 [65.8, 82.9] | 1.15 [1.02, 1.29] | 106.8 [93.0, 126.1] | 1.66 [1.45, 1.96] | 3040 [2692, 3410] | 3688 [3322, 4090] |
| P10 | 62† | 44.1 [33.4, 53.4] | 0.69 [0.52, 0.83] | 78.7 [68.6, 89.0] | 1.22 [1.07, 1.38] | 3235 [2913, 3576] | 4012 [3645, 4401] |
| P11 | 62† | 49.6 [41.7, 57.1] | 0.77 [0.65, 0.89] | 97.5 [86.1, 112.0] | 1.52 [1.34, 1.74] | 2930 [2736, 3138] | 3299 [3153, 3451] |
| P12 | 62† | 79.8 [70.3, 90.9] | 1.24 [1.09, 1.41] | 113.6 [100.3, 134.9] | 1.77 [1.56, 2.10] | 2868 [2689, 3040] | 2696 [2577, 2818] |
| P13 | 59† | n.b. | — | n.b. | — | 3124 [2898, 3354] | 3738 [3063, 4857] |
| P14 | 62 | n.b. | — | — | — | 7460 [6360, 9185] | — |
| P15 | 62 | n.b. | — | — | — | 3786 [3428, 4137] | — |
| P16 | 62 | n.b. | — | — | — | 2948 [2664, 3232] | — |

*Note.* Threshold exclusion criterion: fitted 62.5% correct threshold not bracketed by the tested disparity range (0 to 110 %IPD static; 0 to 137.5 %IPD motion, extending to 139.7 %IPD for P6, P7 and P9–P12 and to 146.8 %IPD for P13, whose motion stimuli were rendered at 63 mm). Reaction time values are mean correct-trial reaction time, pooled across disparity levels and excluding control trials, with bootstrap 95% confidence intervals (Section 2.7). Thresholds are maximum-likelihood estimates with parametric-bootstrap 95% confidence intervals (Section 2.7). Angular values are the maximum within-object disparity, averaged across the four objects and computed from the rendering geometry of each session (Section 2.4). IPD is the measured interpupillary distance; † motion stimuli rendered at 63 mm (Section 2.5). n.b., threshold not bracketed by the tested range; —, not tested. The paired threshold comparison (Section 3.4, *n* = 11) uses only observers with an included threshold in both conditions; the paired reaction time comparison (*n* = 13) uses all observers with valid reaction time data in both conditions, since reaction time does not depend on the psychometric fit converging. P14 to P16 were not invited to the motion sessions because of their poor performance in the static condition (Section 4.4). Static sessions always preceded motion sessions. Observers M2, M3 and M4 of the monocular control (Figure 6) are P15, P12 and P16.

Thresholds could not be reliably estimated for four static condition fits and two motion condition fits, in each case because the fitted 62.5% correct criterion fell outside the range of disparities tested (see Section 2.7 for the pre-specified exclusion rule). This left 12 observers with a usable static threshold, 11 with a usable motion threshold, and 11 observers with a usable threshold in both conditions, who form the paired threshold comparison in Section 3.4. Reaction time analyses did not depend on the psychometric fit converging and therefore included all 13 observers who completed both conditions. Full per-observer coverage is given in Table 1.

### 2.2 Apparatus and stimulus display

Stimuli were presented on a VPixx PROPixx projection system (VPixx Technologies, Saint-Bruno, Canada) running at a refresh rate of 120 Hz. Stereoscopic separation of the left and right eye images was achieved using an active circular polarisation modulator in combination with passive circular polarised glasses worn by the observers. The PROPixx system provides accurate, low-crosstalk stereoscopic presentation and has been validated for vision research, yielding stereo thresholds comparable to those obtained with other established stereoscopic display technologies (Read et al., 2020); crosstalk in stereoscopic displays used for vision research can otherwise be a substantial and often unmeasured source of artefact (Baker, Kaestner, & Gouws, 2016). Display timing was verified at the start of every session, and the experiment aborted if the measured refresh rate deviated from 120 Hz. Stimuli were programmed in MATLAB using the Psychtoolbox (Brainard, 1997; Pelli, 1997) together with the Datapixx/VPixx control libraries. Observers viewed the display binocularly from a fixed distance of 90 cm in a darkened room, with head position stabilised by a chin and forehead rest.

Responses were collected via a VPixx RESPONSEPixx button box, with the four buttons mapped to the four stimulus quadrants; the digital input log of the response device was used to obtain response timestamps synchronised to the display hardware clock, providing precise reaction time measurements. Auditory feedback was delivered after every response through the PsychPortAudio interface: a brief 900 Hz sine tone signalled a correct response and a low-frequency buzzer signalled an incorrect response.

### 2.3 Stimuli

Stimuli were photorealistic renderings of four everyday objects, an apple, a bulb of garlic, a lemon, and a nectarine, generated from high-resolution laser scans of the real objects (Scarfe & Glennerster, 2019). For each object, separate left and right eye images were rendered to produce stereoscopic image pairs. Images were composited onto a uniform grey background (RGB 123, 123, 123) at load time and displayed at a fixed scale, so that each rendered image subtended 15.8° horizontally. On each trial, four objects were presented simultaneously, one centred in each quadrant of the display, with a central white fixation dot (0.14° in diameter) on the grey background (Figure 1). Each object was approximately 4.3° across, and its centre lay 7.7° from the fixation dot.

**Figure 1.**
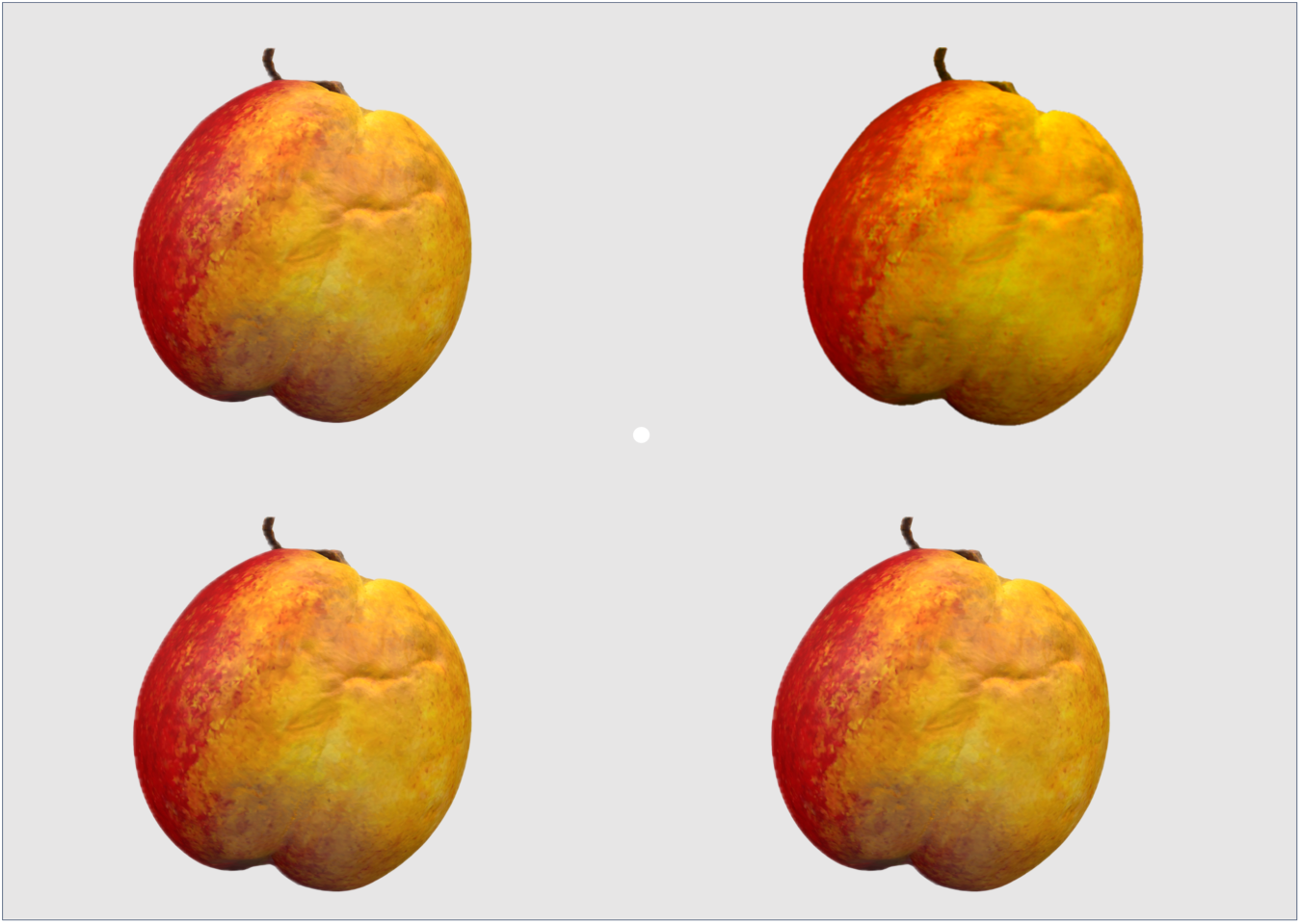
Stimuli and trial display. Example of a photorealistic object (nectarine) rendered from a high-resolution laser scan, showing the shading and surface texture preserved by the scan, schematically showing the four-quadrant binocular display seen by the observer. On each trial one of the four objects, the target, carried added binocular disparity while the three distractor objects carried none; the observer’s task was to report the quadrant containing the target object.

#### Image generation

Stereoscopic image pairs were generated offline in Blender 3.01 (Blender Foundation) using the real-time rasterising renderer. The three-dimensional object models, obtained as laser-scanned meshes with per-vertex colour, were rendered at a resolution of 1920 × 1080 pixels. Each object was illuminated by a fixed multi-source lighting rig (key, fill, rim, and bottom area lights, with intensities and sizes scaled to the camera distance) to produce consistent, naturalistic shading and to reveal the surface texture captured in the scan. Soft shadows and ambient occlusion were enabled. The object was automatically framed to a fixed proportion of the image and rendered against a uniform background. The virtual cameras used a 50 mm lens on a 36 mm sensor.

Left and right eye views were produced by a pair of virtual cameras separated horizontally about the central viewpoint, with a shifted frustum (off-axis) configuration converged at the object centre. The separation between the cameras (the effective stereo baseline, b_eff_) was set as a fraction of the individual observer’s interpupillary distance: b_eff_ = IPD × (%IPD / 100) × *g*. Here %IPD, which we call the rendering level, is the nominal camera separation expressed as a percentage of the observer’s interpupillary distance, and *g* is a small multiplicative gain (0.90 or 1.10) used to sample two values around each rendering level (Section 2.5).

### 2.4 Conversion to angular units

The maximum binocular disparity within each object was converted to angular units in two stages, because the angle subtended at the rendering camera is not the angle subtended at the observer’s eye. Reporting the former as though it were retinal disparity would overstate the latter by a factor of approximately 2.6x under the present display geometry.

First, the horizontal disparity present in the rendered image pair was obtained from the scene geometry as *δ_PX_* = *b_eff_* · (1/*Z_S_* − 1/*Z_C_*) · *f_PX_*, where *b_eff_* is the effective camera baseline for that trial, *Z_C_* the convergence distance, *Z_S_* the distance to the nearest surface point of the object and *f_PX_* = (lens / sensor) × *R_X_* = 2666.7 px the focal length in pixels. *Z_C_* was set at the object centre. *Z_S_* was taken as the first surface point on the ray from the camera through the object centre. *δ_PX_* is the maximum binocular disparity present within the object, between this point and the convergence plane. The maximum is reported because the observer’s task was to detect that one object carried three-dimensional structure while the others were flat, so the relevant signal is the depth range within the object.

Because the images depend only on the ratios between the camera baseline, the object size and the camera distance, the absolute scale of the rendered scene is arbitrary, and all scene lengths are given here in units of the observer’s interpupillary distance (IPD). The camera separation in units of IPD equals the rendering level divided by 100, for example 0.4 IPD at 40 %IPD and 1 IPD at 100 %IPD (Section 2.5), with the two cameras displaced symmetrically about the central viewpoint. Each object was scaled to a common bounding box diagonal of 33.3 IPD, and the camera was placed at the distance that framed it in the image, giving convergence distances Z_C_ of 96.17 IPD (apple), 97.67 IPD (lemon), 87.18 IPD (nectarine) and 84.10 IPD (garlic), and nearest surface distances Z_S_ of 86.45, 88.56, 78.50 and 75.97 IPD respectively. These values are for a 63 mm interpupillary distance; because the scene was the same for every observer while the baseline followed each observer’s interpupillary distance, scene lengths expressed in IPD scale with 63 mm / IPD. For example, for the apple at 100 %IPD, δ_PX_ = 1 × (1/86.45 − 1/96.17) × 2666.7 = 3.12 px.

Second, this disparity was converted to retinal disparity using the display geometry. Each rendered pixel was drawn at 0.59 of its native size (the scale factor applied to the loaded texture), on a projected image 424 mm wide at 1920 px horizontal resolution, viewed from 900 mm. One rendered pixel therefore corresponds to 0.59 × (424/1920) = 0.1303 mm on the screen, subtending 0.4977 arcmin at the eye. Because the scale factor multiplies the texture’s native pixel dimensions, this conversion does not depend on the saved image size. In the example above, 3.12 px therefore corresponds to 1.55 arcmin (0.026°) at the eye. Across the four objects, a rendering level of 100 %IPD produced a maximum within-object disparity of 1.4–1.7 arcmin (0.023–0.028°). Disparities smaller than one screen pixel (0.84 arcmin) were carried by anti-aliasing in the rendering and by luminance (bilinear) interpolation when the scaled texture was drawn.

Relation to depth at the viewing distance. The rendering geometry was not orthostereoscopic. Because each image was displayed at 0.59 of its native size, angles at the eye were about 2.6 times smaller than the corresponding angles at the rendering camera, and the cameras’ convergence point was displayed in the plane of the screen, so the observer’s vergence and accommodation were both at the viewing distance of 900 mm. The rendered camera distance (84–98 IPD) therefore has no counterpart in the stimulus and is not specified by any cue available to the observer. At the viewing distance, each displayed object subtended about 4.3°, the angular size of a 68 mm object at 900 mm, and at 100 %IPD its maximum within-object disparity (1.4–1.7 arcmin) is the disparity produced, for a 63 mm interpupillary distance, by a binocular depth of 5–6 mm between the object’s nearest point and its centre. In respect of binocular depth, the displayed objects were therefore much flatter than real fruit of the same angular size, for which this depth would be about 34 mm; at the mean thresholds (0.89 arcmin static, 1.48 arcmin motion) the equivalent depth was about 3.3 mm and 5.5 mm. The %IPD values describe the camera geometry used for rendering, and the angular values describe the stimulus delivered to the eye.

Two consequences of this scheme should be noted. Because disparity scales with the observer’s interpupillary distance by construction, a given nominal %IPD corresponds to a different absolute disparity in different observers; each observer’s threshold was therefore converted using their own measured interpupillary distance, and the resulting angular values averaged. Also, because the four object types differ in depth profile, the disparity produced by a given baseline differs slightly between them by approximately 9% (coefficient of variation across objects, 0.087 at every interpupillary distance tested), so a given rendering level corresponds to a narrow range of angular disparities rather than to a single value. Angular thresholds below are means across the four objects.

### 2.5 Disparity parameterisation

The magnitude of the added binocular depth in the target object was defined relative to each observer’s interpupillary distance and expressed as a percentage of that distance (%IPD; the rendering level), because the rendered stereo baseline was set as a fraction of the assumed interocular separation. The effective camera baseline was computed as b_eff_ = IPD × (%IPD / 100) × g, where g is the disparity gain (0.90 or 1.10; see Section 2.3 and below), so that at 100 %IPD the cameras were separated by exactly the observer’s interpupillary distance (Section 2.4). A value of 0 %IPD therefore corresponded to identical left and right eye images, and increasing %IPD corresponded to rendering the object with a progressively larger effective interocular separation, thereby increasing the binocular depth signal (Figure 2). Stereo pairs were pre-rendered at each required level for each observer’s measured interpupillary distance, so that the disparity content of every stimulus was fixed by the rendering geometry rather than by run-time image manipulation. For seven observers (P6, P7 and P9–P13), the motion-condition stimuli were rendered at an interpupillary distance of 63 mm rather than at the observer’s measured value (62 mm for P6, P7 and P9–P12; 59 mm for P13). For these observers, motion-condition disparities are expressed as a percentage of the measured interpupillary distance (nominal %IPD × 63 / measured interpupillary distance), so that %IPD has the same meaning in both conditions, and angular disparities were computed from the geometry actually rendered (Section 2.4).

**Figure 2.**
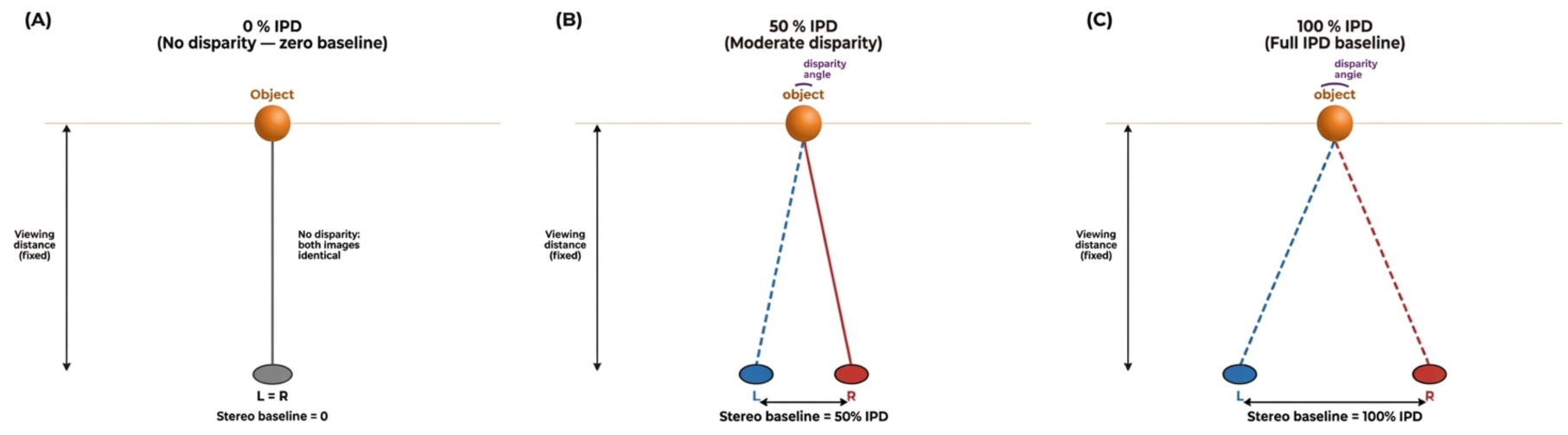
Disparity parameterisation in %IPD. Rendering geometry for three representative levels. At 0 %IPD the stereo baseline is zero and the left and right eye images are identical, so the object carries no binocular disparity. At 50 %IPD and 100 %IPD the virtual cameras are separated by the corresponding fraction of the observer’s interpupillary distance, producing progressively larger disparity angles at the object. Viewing distance is fixed throughout.

To enable fine estimation of the detection threshold, each baseline level was additionally modulated by a small multiplicative disparity gain producing two flanking values around the nominal level (gain factors of 0.90 and 1.10). In the analysis the stimulus level on each trial was therefore taken as the *effective* %IPD, defined as the nominal baseline multiplied by its gain factor, giving a finely sampled disparity axis around each nominal level (Figure 3).

**Figure 3.**
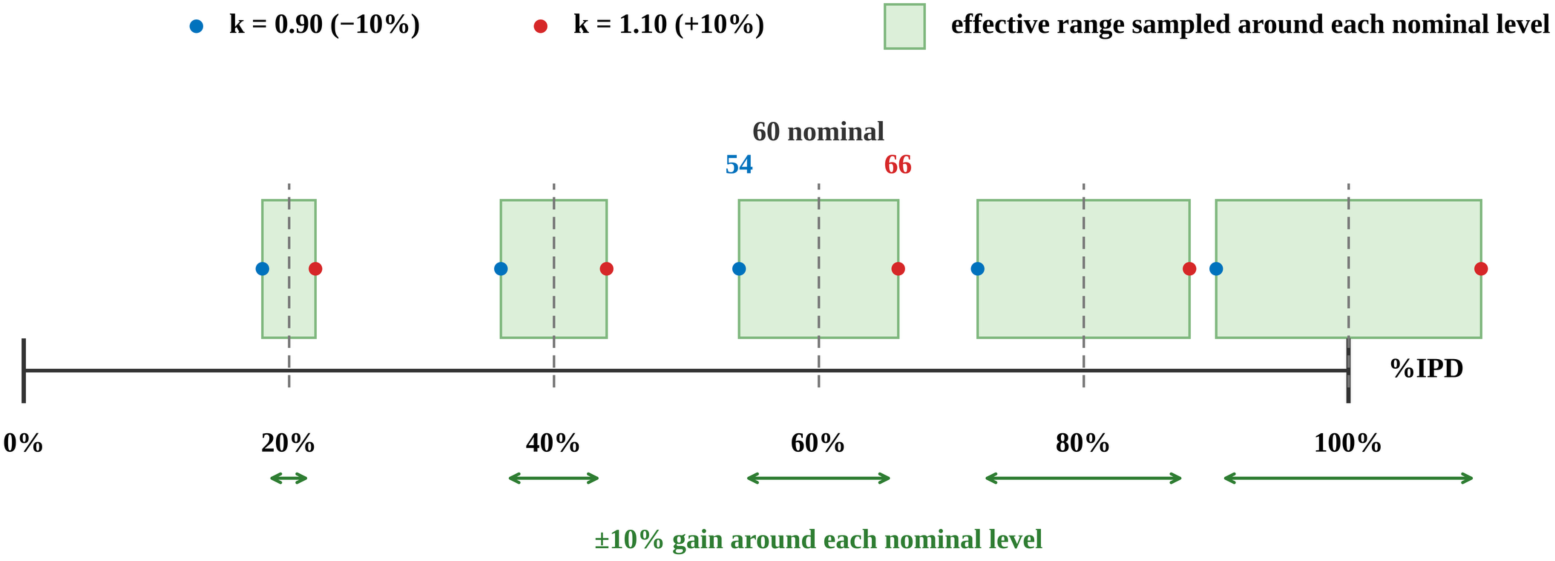
Disparity gain sampling. Each nominal %IPD level was sampled at two effective values by applying multiplicative gains of 0.90 and 1.10. For example, a nominal level of 60 %IPD produced effective values of 54 and 66 %IPD. This fine sampling around each nominal level provides the resolution needed for stable psychometric fitting.

**Figure 4.**
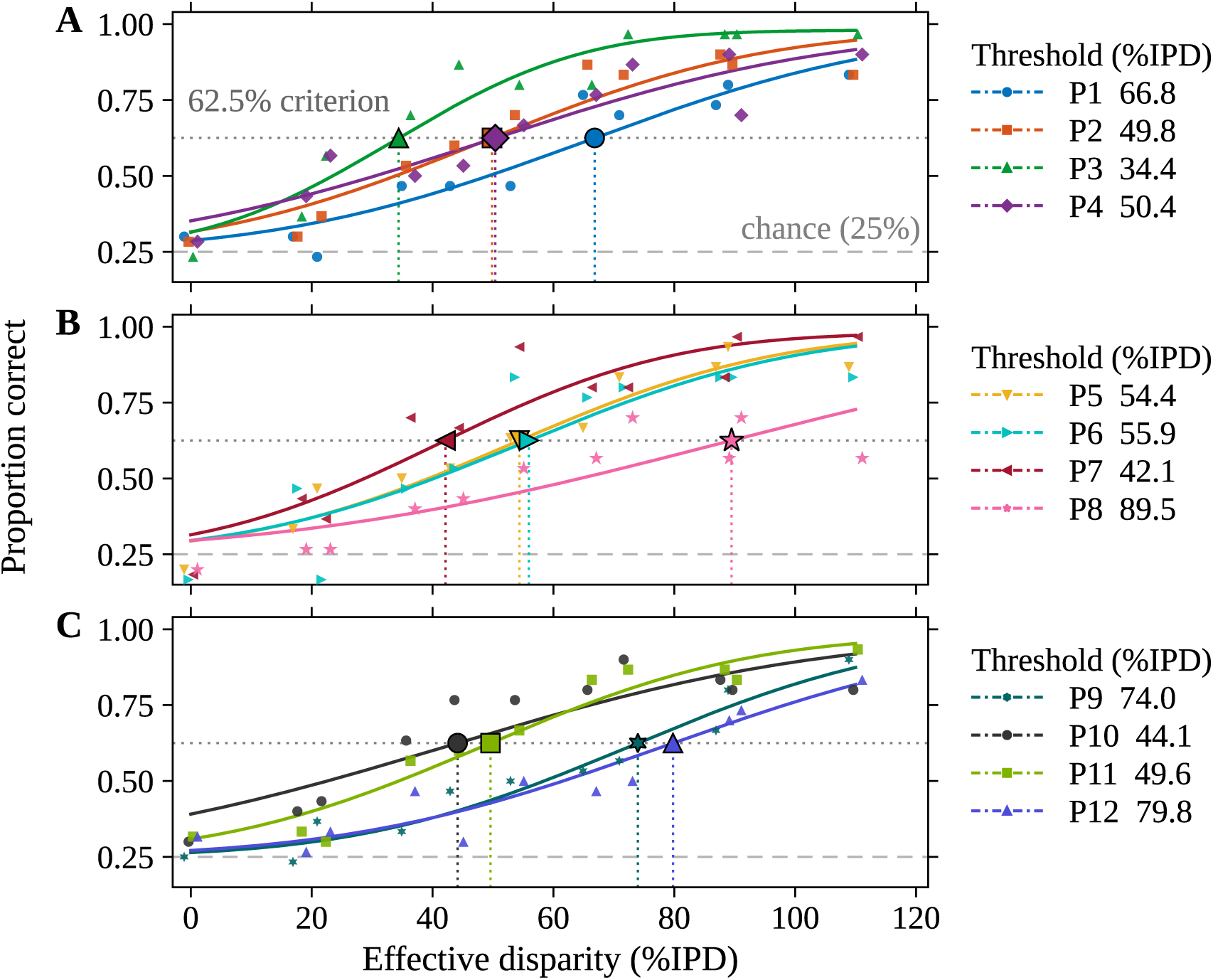
Psychometric functions for the static condition. Proportion correct is plotted against effective disparity (%IPD) for the 12 observers whose threshold fell within the tested range (A: P1–P4; B: P5–P8; C: P9–P12). Points show observed proportion correct at each effective level, based on 30 trials per level (60 at 0 %IPD), offset horizontally by up to ±1.1 %IPD for visibility. Curves show maximum-likelihood cumulative Gaussian fits with guess rate fixed at 0.25 and lapse rate at 0.02. Large outlined markers with dotted drop lines mark each observer’s threshold (62.5% correct), and the legend gives each threshold value. The dashed line marks chance (25%) and the dotted line the 62.5% criterion. P13–P16 are not shown because their fitted thresholds fell outside the tested range (see Table 1). Group mean threshold: 57.6 %IPD (0.89 arcmin). Each observer has the same colour and marker in Figs 4, 5 and 8.

**Figure 5.**
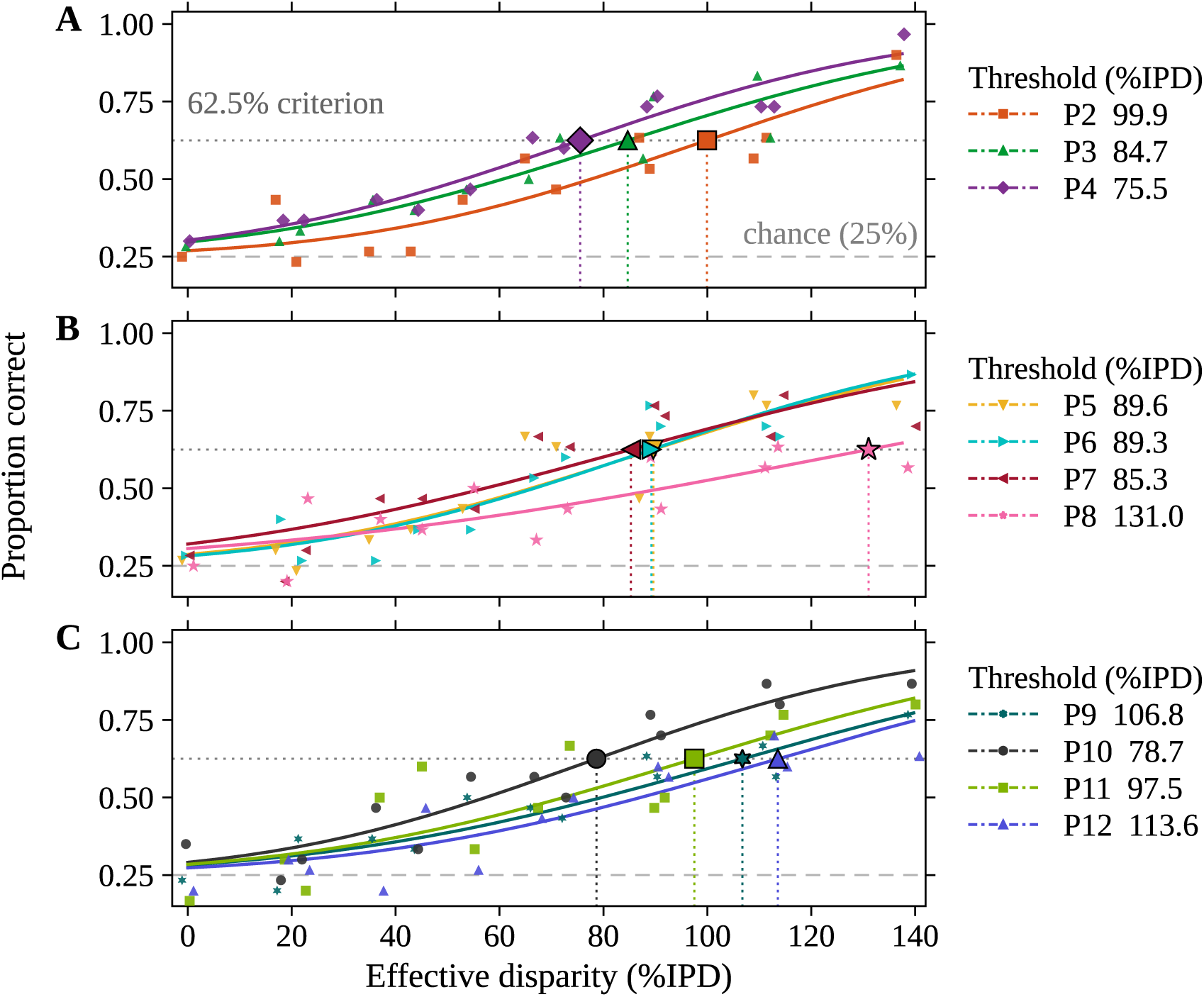
Psychometric functions for the motion condition, in which the objects rotated. Conventions are as in Figure 4, for the 11 observers whose threshold fell within the tested range (A: P2–P4; B: P5–P8; C: P9–P12). P1 and P13 are not shown because their fitted thresholds fell outside the tested range, and P14–P16 were not tested in this condition (see Table 1). Each point is based on 30 trials (60 at 0 %IPD). The tested range extends to 137.5 %IPD because the highest nominal level (125 %IPD) was sampled with a gain of 1.10. Group mean threshold: 95.6 %IPD (1.48 arcmin).

### 2.6 Task and procedure

Observers performed a spatial four-alternative forced-choice odd-one-out task (Jäkel & Wichmann, 2006). The four-quadrant locations were occupied by four renderings of the same object; on a given trial, three distractor objects were presented with no added binocular depth, and one target object, chosen at random with equal probability across the four locations, carried the added disparity signal. The observer reported the location of the target object by pressing the corresponding button. Chance performance was therefore 25% correct. Observers were instructed to fixate the central dot and, once it disappeared, to find the object that differed from the other three and report its location as quickly and accurately as possible.

Each trial proceeded through a fixed sequence of epochs: a fixation period (0.5 s) during which only the central fixation dot was visible; a pre-target presentation period (1.0 s) during which all four objects were displayed without binocular disparity, the same image being shown to both eyes; the target period (2.0 s), during which one of the four objects carried the added disparity signal, after which it reverted to zero disparity while all four objects remained visible. Finally, an inter-trial interval (0.5 s) was enforced. Observers could respond at any time from target onset, and reaction time was measured from target onset. On every trial, the three distractor objects were the left-eye image of the stereo pair, shown to both eyes. Auditory feedback followed each response.

#### Motion condition

In a separate set of sessions, the four objects rotated about a vertical axis to provide a structure-from-motion cue in addition to disparity, shading, outline shape and other cues. For each object and disparity level, a 24-frame stereoscopic sequence was rendered, depicting rotation about the vertical axis. Each sequence lasted 0.2 s and comprised 24 frames, presented at 120 frames per second with a rotation speed of 180°/s. The sequence therefore spanned 36° of rotation. During presentation, the sequence was played back frame-locked to the 120 Hz display in a continuous forward and reverse loop, so that the object oscillated smoothly through the rotation range. The camera rig remained fixed in world coordinates, so the motion arose from the object turning relative to a stationary observer. During the pre-target period, the rotating objects were presented without added binocular disparity; at target onset, the target object additionally acquired the disparity signal while continuing to rotate.

#### Monocular control

To verify that the target object was detected by its binocular disparity, and not by any other feature of the trial, a control was run in four observers (M1–M4; Section 3.2). Control trials were interleaved with the main trials of a static session: for M2, M3 and M4 (P15, P12 and P16) this was their static session of the main experiment, and for M1 a separate static session run for the control. Control trials used the same task, objects and trial sequence, but the target object was shown with its left-eye image in both eyes, so that it carried no binocular disparity. The three distractor objects were the same as in the main experiment. Control trials were therefore not assigned a %IPD level. If observers could locate the target from anything other than its binocular disparity, performance on control trials should have remained above chance; performance at chance would indicate that the task could be performed only when binocular disparity was present. Controls of this kind are important whenever fully shaded stimuli are used to probe stereopsis, since monocular artefacts are a recognised confound in stereo testing (Serrano-Pedraza, Vancleef, & Read, 2016).

#### Design

The two main conditions were run in separate sessions. For static objects, six baseline levels were tested (0, 20, 40, 60, 80, 100 %IPD), with 30 trials per gain value, giving 360 trials per participant. For moving objects, seven baseline levels were tested (0, 20, 40, 60, 80, 100, 125 %IPD), giving 420 trials per participant. Each session was preceded by a short block of practice trials at a high disparity level. Static sessions were always run before motion sessions. A further feature of the design was that, because the task required detection of which object differed, any global change applied equally to all four objects could not by itself signal the target object; only a difference between the target object and the three distractor objects was informative.

### 2.7 Data analysis

Only trials from the main experimental phase were analysed. For each session, the two disparity gain values were incorporated by expressing the stimulus level on each trial as an effective %IPD. For each observer and condition, the proportion of correct responses was computed at each unique effective level (the two gain values at each nominal level were treated as separate stimulus levels, not averaged) and fitted with a cumulative Gaussian psychometric function of the form

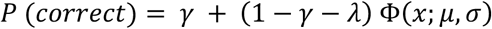

where Φ is the cumulative normal distribution, the guess rate was fixed at γ = 0.25 (appropriate for the four-alternative design), and the lapse rate was fixed at λ = 0.02 (Wichmann & Hill, 2001a). The location (μ) and slope (σ, the standard deviation of the cumulative Gaussian) parameters were estimated by maximum likelihood, treating the number of correct responses at each effective level as binomially distributed (Wichmann & Hill, 2001a). Goodness of fit was assessed from the deviance of each fit, compared with the distribution of deviances obtained by refitting 1,000 data sets simulated from the fitted function (Wichmann & Hill, 2001a). The detection threshold was defined as the effective %IPD yielding 62.5% correct, the midpoint between chance and the upper asymptote, and was calculated directly from the fitted function. Ninety-five per cent confidence intervals on each observer’s threshold were obtained by parametric bootstrap (2,000 resamples; Wichmann & Hill, 2001b): binomial responses were simulated from the fitted function at the tested levels and trial numbers, refitted with the same procedure, and the 2.5th and 97.5th percentiles of the resulting thresholds were taken.

Observers whose fitted threshold fell outside the range of tested disparities, that is where the 62.5% correct criterion was not bracketed by the data and the reported value would therefore be an extrapolation, were excluded from the group statistics under this pre-specified rule.

To test the effect of motion within each observer, the static and motion data of that observer were fitted jointly with two nested models (Kingdom & Prins, 2016): a fuller model, in which the two conditions had separate thresholds, and a lesser model, in which they shared a single threshold. Both models allowed a separate slope for each condition. The models were compared by a likelihood-ratio test, with p values obtained by Monte Carlo simulation: both models were refitted to 500 data sets generated from the lesser model, and p was the proportion of simulated likelihood ratios at least as large as the observed one (Scarfe & Glennerster, 2014).

Reaction time was measured from target onset to the response and analysed separately for correct and incorrect trials. For each observer, a 95% confidence interval on mean correct-trial reaction time was obtained by bootstrap resampling of that observer’s correct trials (2,000 resamples). Control trials were excluded from all reaction time analyses. Differences between the static and motion conditions in detection threshold and in mean correct-trial reaction time were assessed with paired-samples *t-*tests, matching observers across conditions, with Wilcoxon signed-rank tests as nonparametric companions. Because reaction-time distributions are positively skewed, the reaction time analyses were repeated using each observer’s median correct-trial reaction time. Control performance was compared against the 25% chance baseline using exact binomial tests, with Clopper–Pearson confidence intervals. Psychometric fitting and model comparison were performed in Python (SciPy); other analyses were performed in MATLAB.

## 3. Results

All observers were able to perform the odd-one-out task, indicating that binocular disparity carried in photorealistic objects supports reliable depth detection. For each observer, the proportion of correct responses increased monotonically with effective disparity, and individual psychometric functions were well described by a cumulative Gaussian.

### 3.1 Disparity detection with photorealistic objects

For static objects, 12 of 16 observers yielded a threshold estimate within the tested range of disparities. Converting each observer’s threshold at their own rendered interpupillary distance (Section 2.4), the mean detection threshold in this group was a maximum within-object disparity of 0.89 arcmin (*SD* = 0.24; 0.0149°), with individual thresholds ranging from 0.54 to 1.30 arcmin. This corresponds to a mean rendering level of 57.58 %IPD (*SD* = 16.54, range 34.40 to 89.49).

For moving objects, 11 of 13 observers who completed this condition yielded a threshold within range. The mean threshold was 1.48 arcmin (*SD* = 0.23; 0.0247°), with individual thresholds ranging from 1.19 to 1.91 arcmin, corresponding to a mean rendering level of 95.62 %IPD (*SD* = 16.48, range 75.53 to 131.04). Without the exclusion (*n* = 13), the mean rendering level at threshold was 107.50 %IPD (*SD* = 34.61). The cumulative Gaussian described the data well: for 26 of the 29 fits across both conditions the deviance lay within the range expected from simulated data (p > 0.05). The three exceptions (static condition, P6 and P7; motion condition, P11) all had thresholds well within the tested range.

### 3.2 Detection depends on binocular disparity

Performance in the main experiment covaried with the disparity added to the target object, but covariation alone does not establish that observers were using binocular disparity. Rendering an object with a larger effective interocular separation changes the left and right eye images in ways that are not confined to their binocular relationship, each monocular image is itself drawn from a slightly displaced viewpoint, and any residual cue that scaled with disparity level, such as a shift in the visible silhouette, in the pattern of shading, or in the position of a specular highlight, could in principle have supported detection without binocular fusion. This concern is specific to photorealistic stimuli. It does not arise for random dot stereograms, in which the three-dimensional structure is invisible to either eye alone, and it is the price of using objects that carry a full complement of pictorial cues.

The monocular control removed the binocular disparity of the target object by presenting its left-eye image to both eyes. If the observers could perform the task on the basis of the slight change in object viewpoint due to the slight difference in left-eye camera axis, we might expect some ability to pick the odd stimulus out of the four. Performance on control trials was at chance: pooled across observers, accuracy was 26.2% (95% CI [19.6, 33.8], *N* = 160 trials), not different from the 25% expected by guessing, and no individual observer differed from chance (Figure 6A). The upper limit of this interval is informative: the data exclude accuracy above roughly 34%, that is, more than about 9 percentage points above chance, so the absence of an effect reflects adequate sensitivity rather than an uninformative test. In the same observers, binocular accuracy rose monotonically from chance at 0 %IPD to approximately 58% at 100 %IPD, whereas accuracy on control trials, which carried no binocular disparity, remained at chance (Figure 6B).

**Figure 6.**
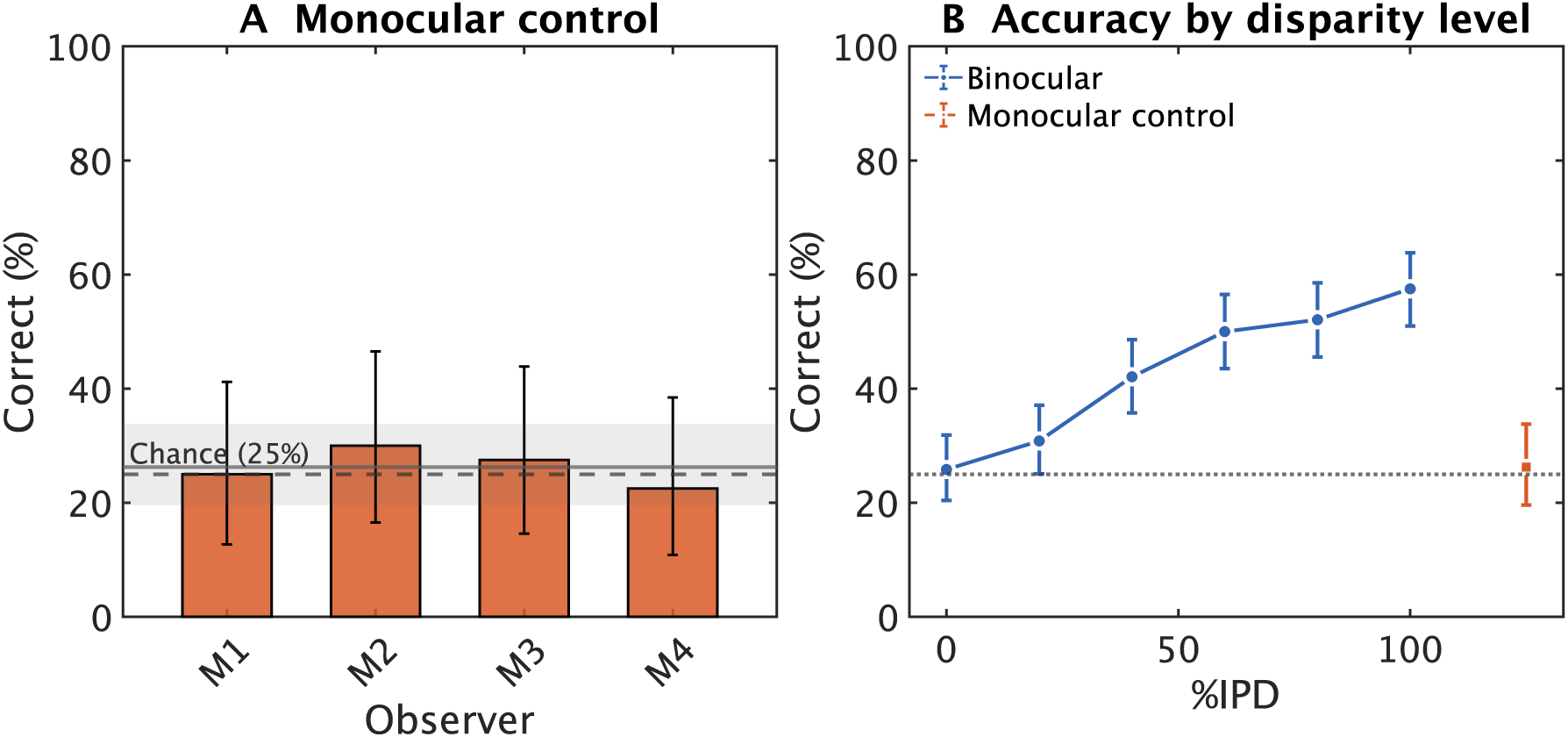
Detection requires binocular disparity. **(A)** Accuracy on monocular control trials for each observer (M1–M4), in which the left-eye image of the target object was presented to both eyes. M2, M3 and M4 are P15, P12 and P16; M1 is a further control session run separately from the main experiment. Bars indicate individual observer accuracy; error bars represent exact binomial 95% confidence intervals; grey line and shaded band represent the pooled estimate; horizontal dashed line indicates chance performance (25%). **(B)** Average binocular accuracy as a function of disparity (blue data points) and the monocular control (red data point) for the four observers who completed control trials (M1–M4), pooled across observers without the threshold-range exclusion applied in Figs 4–5; two of these observers had no threshold within the tested range, which accounts for the lower accuracy compared with Fig 4.

Because the two trial types were interleaved within session, drew on the same objects and the same rendered images, and differed only in whether the two eyes received different views, this comparison isolates binocular disparity as the effective variable. Detection in the main experiment therefore depended on the binocular disparity of the target object.

### 3.3 Reaction time decreases with disparity

Reaction times on correct trials provided an independent measure of disparity sensitivity. In both conditions, correct-trial reaction time decreased systematically as the added disparity increased: larger, more easily detected disparities were associated with faster responses. In the static condition, mean correct-trial reaction time fell from 4112 ms (*SD* = 1720) at 0 %IPD to 2461 ms (*SD* = 1101) at 100 %IPD (Figure 7A). The motion condition showed the same monotonic decline, from 4086 ms (*SD* = 1809) at 0 %IPD to 2574 ms (*SD* = 518) at 125 %IPD (Figure 7B). The narrowing of the standard deviation with increasing disparity is itself informative: at low disparities, observers differ widely in how long they deliberate before responding, whereas at high disparities, where the target is readily visible, they converge on a similar response latency.

**Figure 7.**
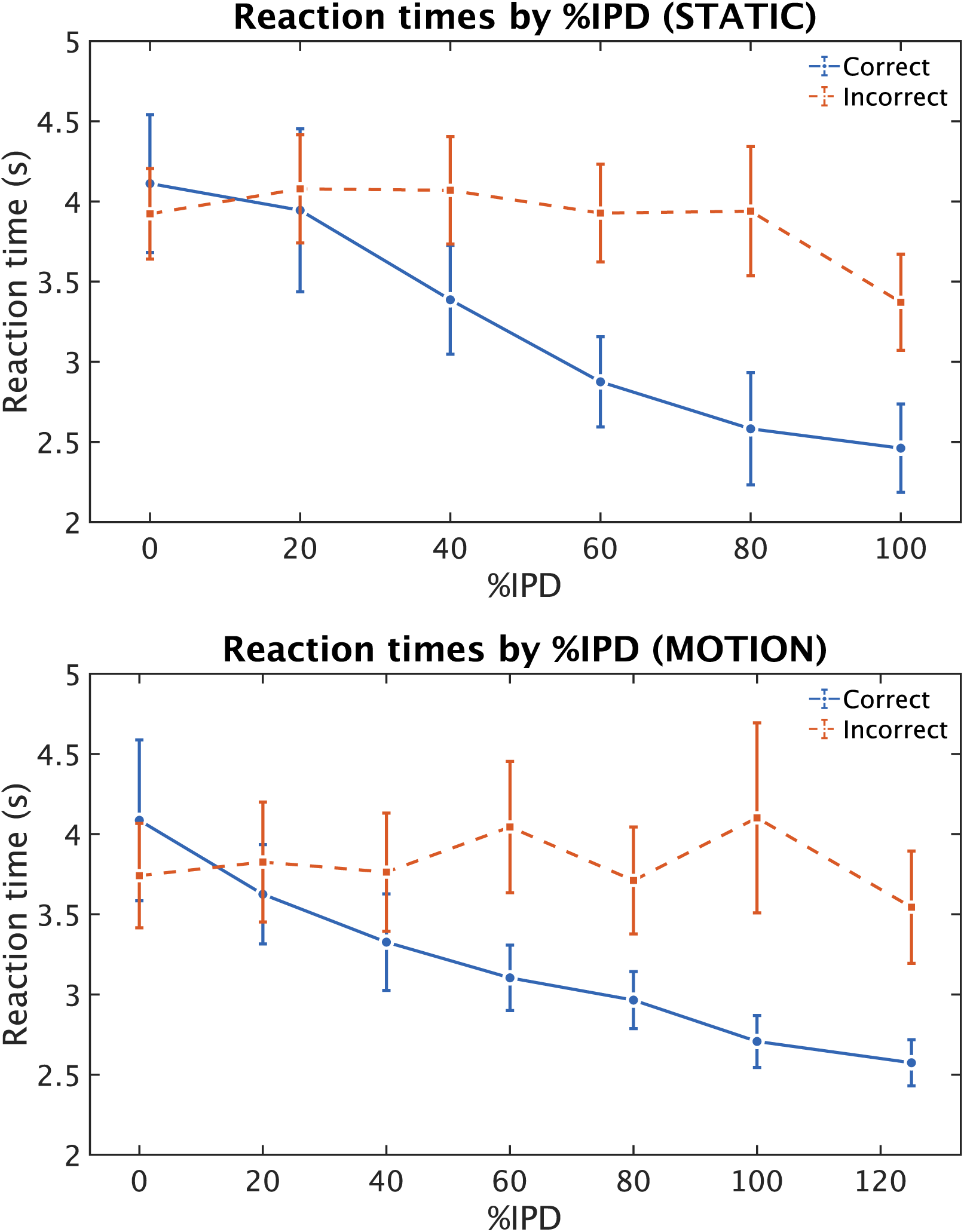
Reaction time as a function of disparity level. Pooled mean reaction time for correct (blue, solid) and incorrect (orange, dashed) trials in the static condition (top) and the motion condition (bottom). Correct-trial reaction time decreases monotonically with increasing disparity in both conditions. Incorrect-trial reaction time shows no significant modulation in the motion condition and only a smaller residual decrease in the static condition. Error bars indicate the standard error of the mean.

**Figure 8.**
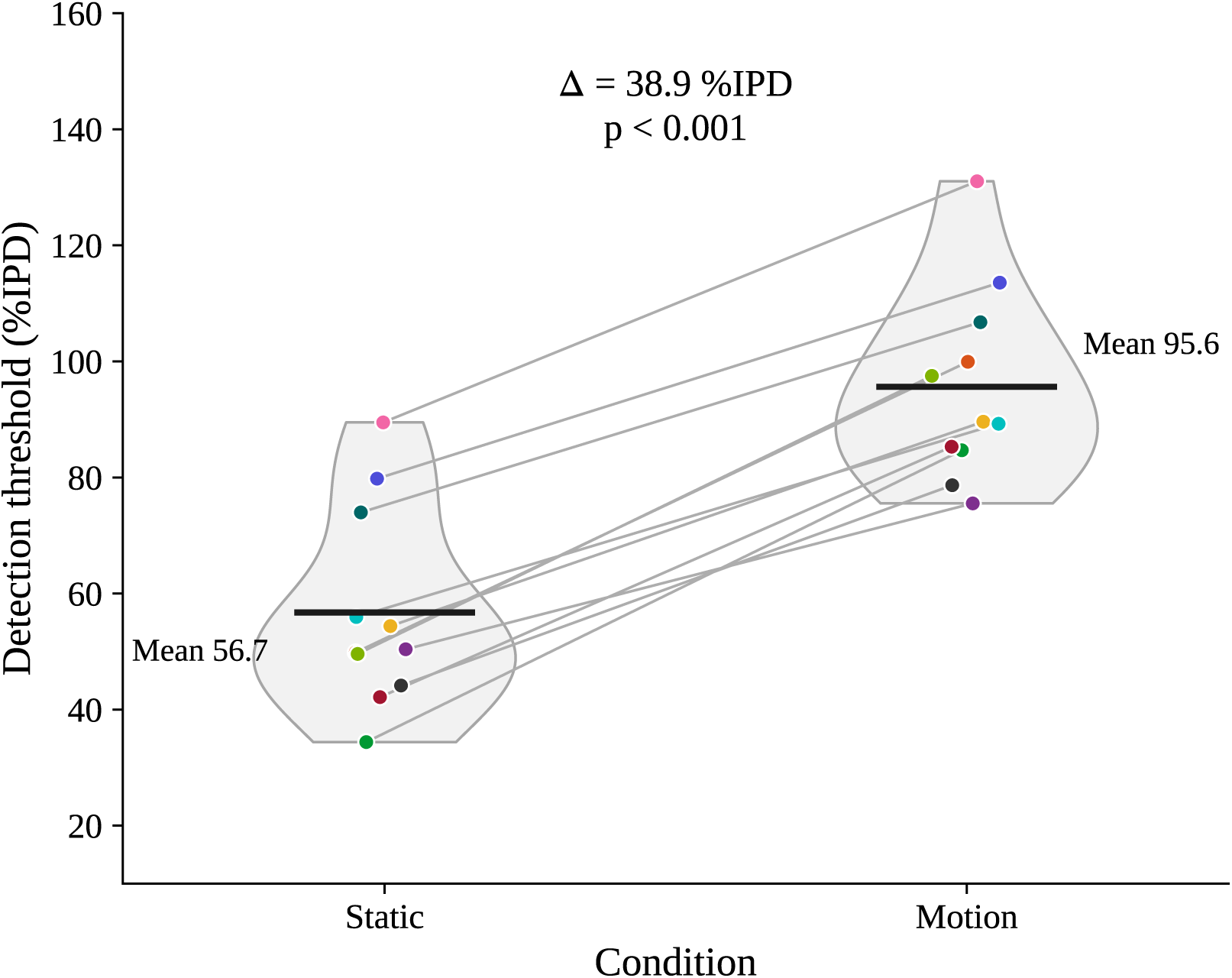
Static versus motion thresholds for the 11 observers with a usable threshold in both conditions. Coloured points: individual observers (colours as Figs 4–5), joined by grey lines; violins: density of thresholds per condition; black bars: group means (56.7 and 95.6 %IPD). Every observer is higher under motion (mean difference 38.9 %IPD, 95% CI [33.3, 44.4]; paired t(10) = 15.66, p < 0.001).

A repeated-measures analysis of variance confirmed this effect of disparity level on correct-trial reaction time in both the static condition, *F*(5, 75) = 12.17, *p* < 0.001, partial η² = 0.448, and the motion condition, *F*(6, 72) = 9.15, *p* < 0.001, partial η² = 0.433. Sphericity was strongly violated in both cases (Greenhouse–Geisser ε = 0.409 and 0.348 respectively), reflecting the larger variance in reaction time at low disparity; the effect remained significant after correction (both *p* < 0.001). Because the prediction is specifically a monotonic decrease rather than any difference among levels, we also tested the linear trend directly, which was significantly negative in both conditions (static, *t*(15) = −4.70, *p* < 0.001; motion, *t*(12) = −4.56, *p* < 0.001). The linear trend was also significant when each observer’s median correct-trial reaction time was used in place of the mean (static, t(15) = −3.92, p = 0.001; motion, t(12) = −3.72, p = 0.003).

This decrease in reaction time with disparity was substantially stronger on correct than on incorrect trials, though the dissociation was not absolute. In the motion condition, incorrect-trial reaction time showed no effect of disparity, *F*(6, 72) = 1.10, *p* = 0.372, partial η² = 0.084, exactly the pattern expected if errors arise from trials on which the signal was not detected and the observer guessed. In the static condition, incorrect-trial reaction time did decline modestly with disparity, *F*(5, 75) = 3.70, *p* = 0.005 (Greenhouse–Geisser corrected *p* = 0.040), but the effect was less than half the size of that on correct trials (partial η² = 0.198 versus 0.448) and is confined to the highest tested disparity. A residual decline on error trials is not unexpected: at high disparities, errors are increasingly likely to be lapses or premature responses rather than genuine failures of detection, and on average these might be faster responses. The important comparison is the magnitude of the two effects. By that measure, reaction time tracks the detection process rather than a generic speeding of all responses. The same inverse relationship between reaction time and task difficulty is observed in macaques performing the four-quadrant depth change detection task (Smith & Parker, 2021), which supports the cross species comparability of the paradigm.

### 3.4 Static versus moving objects

Comparing static and motion thresholds within observer requires restricting the comparison to participants with a usable threshold in both conditions, since four static and two motion fits were excluded as extrapolated, and three observers did not complete the motion condition. Eleven observers met this criterion and form the paired comparison reported here; full per-observer coverage is given in Table 1.

Within this matched sample, detection thresholds were consistently higher for moving objects in comparison with static ones. Mean threshold rose from 56.74 %IPD for static objects to 95.62 %IPD for moving objects, an increase of 38.88 %IPD (95% CI [33.35, 44.42]). The difference was highly reliable, paired *t*-test, *t*(10) = 15.66, *p* < 0.001, and large in magnitude, Cohen’s *d*z = 4.72; a Wilcoxon signed-rank test gave the same conclusion (*p* = 0.001), indicating the result does not depend on the normality assumption of the parametric test. Critically, every one of the 11 paired observers showed a threshold increase under motion, with individual increases ranging from 25.14 to 50.28 %IPD. The effect was uniform in direction across the sample, not driven by a subset of observers. The model comparison (Section 2.7) confirmed this for each observer individually: for all 11 observers, a model with separate static and motion thresholds fitted the data significantly better than a model with a single shared threshold (likelihood ratio 6.6 to 51.9; Monte Carlo p = 0.010 for P8 and p ≤ 0.002 for the other ten; Supplementary Table S9). Expressed in angular units, the mean threshold rose from 0.88 arcmin (0.0146°) for static objects to 1.48 arcmin (0.0247°) for moving objects, an increase of 0.60 arcmin, or 69%.

Correct trial reaction time showed a converging pattern. Across the 13 observers who completed both conditions, mean correct-trial reaction time was 2619 ms for static objects and 3049 ms for moving objects, an increase of 430 ms under motion, paired *t*-test, *t*(12) = 3.15, *p* = 0.008, Cohen’s *d*z = 0.87; the result was the same with median reaction times (t(12) = 3.36, p = 0.006). The effect was consistent in direction across most of the sample but not universal: 11 of 13 observers were slower under motion, while two were faster, by 614 ms and 171 ms respectively. Because static sessions always preceded motion sessions, practice would, if anything, have shortened reaction times in the motion condition; the two reversals may therefore reflect practice or individual variability, and the overall slowing under motion was found despite this ordering. Together, the higher thresholds and longer reaction times indicate that disparity is detected less sensitively and more slowly when it is embedded in a moving, rotating object than when it is carried by a static one.

## 4. Discussion

We measured binocular disparity detection thresholds for photorealistic objects, rendered from high-resolution laser scans, using a spatial four-alternative forced-choice odd-one-out task. Four findings emerged. First, observers reliably detected binocular disparity carried by photorealistic objects, with orderly psychometric functions and well-defined thresholds. Second, removing the binocular disparity of the target object reduced performance to chance, establishing that detection depended on binocular disparity. Third, both accuracy and correct-trial reaction time were modulated by disparity magnitude, while incorrect-trial reaction time showed substantially weaker modulation. Fourth, disparity detection was markedly less efficient for rotating objects than for static ones, thresholds were substantially higher and reaction times longer when structure-from-motion accompanied the disparity signal.

### 4.1 Reliable stereopsis in photorealistic objects

That disparity thresholds could be measured at all with complex, object-rich stimuli is a non-trivial result. Most of what is known about human stereoacuity derives from sparse, abstract stimuli that deliberately strip away monocular form (Julesz, 1971; Cumming & DeAngelis, 2001; Howard & Rogers, 2002; McKee et al., 1990). Photorealistic objects differ on many dimensions at once: they carry shading, texture, occlusion, and recognisable shape, and these monocular cues could in principle have masked the added binocular signal or dominated the depth percept. There were also specific grounds for pessimism. Depth judgements on smoothly curved binocular surfaces are systematically distorted rather than veridical (Hornsey et al., 2016; Hibbard et al., 2012), and observers sample the disparity field non-uniformly when estimating three-dimensional shape (Scarfe & Hibbard, 2013), so it was not obvious in advance that a clean, monotonic psychometric function would be recoverable from a fully shaded, curved object.

Our finding that observers exploit disparity in an orderly, threshold-governed manner indicates that binocular processing remains effective when disparity is embedded in the context of a naturalistic object, consistent with reports that binocular information continues to contribute to perceived depth in stereoscopic photographs of natural scenes (Hibbard et al., 2023; Hornsey et al., 2015) and with the broader view that disparity is integrated with, rather than overridden by, other cues during natural viewing (Landy et al., 1995; Hibbard et al., 2025). It is worth emphasising that the distortions documented for curved surfaces concern the metric accuracy of a suprathreshold depth estimate, whereas the present measure is the detectability of an added binocular signal. The two need not covary, and our results suggest that a depth representation which is sometimes metrically biased can nonetheless support sensitive detection.

The monocular control supports this conclusion. When the binocular disparity of the target object was removed, by presenting its left-eye image to both eyes, performance fell to chance, so detection depended on the difference between the two eyes’ images of the target. Because the control used only the left-eye image, it does not by itself exclude a cue carried by the right-eye image alone; presenting each eye’s image to both eyes, in separate trials, would test this directly. Controls of this kind are important whenever complex, fully shaded stimuli are used to probe stereopsis (Serrano-Pedraza et al., 2016).

The thresholds obtained here, 0.89 arcmin for static objects and 1.48 arcmin for moving objects, are coarser than the finest stereoacuities reported under optimised conditions, which can approach a few seconds of arc (Howard & Rogers, 2002; McKee et al., 1990). This is expected given the differences in task and stimulus: the highest stereoacuities are obtained with high luminance contrast stimuli in the form of a foveally-fixated, optimised target, whereas the present paradigm measured detection of an added disparity on a complex object presented briefly at one of four peripheral locations. The values are also consistent with the wide range of stereo performance documented in population studies of binocular function (Bosten et al., 2015; Coutant & Westheimer, 1993). They are also of the same order as the discrimination thresholds obtained with natural stereo-image patches, roughly 1 to 2 arcmin across pedestal disparities and levels of local depth variation (White & Burge, 2025), although differences in task (spatial four-alternative detection versus two-interval forced-choice discrimination about a pedestal) and threshold criterion preclude a direct numerical comparison. The relevant point for present purposes is not the absolute value but that the measurement is stable and orderly across observers, and obtained with stimuli that approximate natural viewing far more closely than conventional stereo stimuli while remaining quantitatively controllable (Scarfe & Glennerster, 2019).

### 4.2 The cost of motion for disparity detection

The most striking result is that adding rotational movements to the object raised the disparity detection threshold substantially and slowed responses. At first sight this may seem to conflict with classic demonstrations that stereopsis and structure-from-motion combine synergistically: when both cues specify the same three-dimensional shape, the perceptual distortions that each cue produces alone are reduced, and combined-cue shape judgements are more veridical than single-cue judgements (Johnston et al., 1994; Tittle & Braunstein, 1993; Tittle & Perotti, 1997). However, those studies addressed the perceived shape of a clearly suprathreshold object, whereas the present task required detection of a small added disparity in the presence of motion. These are different computational problems, and there are several non-exclusive reasons why motion could impair disparity detection even if motion and disparity cooperate in shape perception.

The first possibility is that the rotation continuously changes the disparity field across the surface of the object, so that the binocular signal an observer must detect fluctuates over the presentation interval; integrating disparity across a changing surface is likely to be noisier than reading a fixed disparity from a static object. A related finding is that disparity-defined objects moving in depth fail to elicit three-dimensional shape constancy (Scarfe & Hibbard, 2006). That study concerned objects translating in depth rather than rotating, but it shows that the visual system does not straightforwardly compensate for a binocular geometry that changes over time. Related evidence comes from natural stereo-images, in which variation in local depth structure acts as a source of stimulus-driven "nuisance" variability that raises disparity thresholds (White & Burge, 2025); a rotating object continuously re-samples its local depth structure, which may impose an analogous cost. If the depth signal is not stabilised across the motion sequence, the effective signal available for detection is correspondingly degraded. A separate issue is that when both stereo and motion signals are near threshold, these cues show summation when presented together (Bradshaw & Rogers, 1996). At first sight this seems in contradiction to the present results, in that here motion is detrimental to stereo thresholds. However, the Bradshaw and Rogers experiment differs in two important ways from the present one. First, the task is different. In the present task, motion is always present but provides no useful information for solving the task, which is to detect which stimulus is different solely on the basis of binocular disparity. Second, the stimuli used by Bradshaw and Rogers are completely smooth in their depth profile and therefore do not contain the local variation of depth structure associated with natural objects and identified by White and Burge (2025).

A second possibility is that the rotating object provides a salient structure-from-motion cue to its three-dimensional shape that is largely independent of the small added disparity; if observers extract object shape from motion alone, the incremental binocular signal becomes comparatively less informative and harder to isolate. Under a reliability-weighted combination scheme, a highly reliable motion-defined shape estimate would receive a large weight and the small disparity increment a correspondingly small one, reducing its influence on the perceptual decision (Landy et al., 1995; Ernst & Banks, 2002). Related to this, the integration of biased sensory estimates does not follow the simple variance-weighted prediction when the component estimates carry systematic error (Scarfe & Hibbard, 2011), which is plausibly the situation here. Another aspect of a reliability-weighted combination scheme as it applies to the stimuli used here is that motion and stereo may be processed at different scales of spatial integration. Arguably, if motion is integrated at a coarser spatial scale, this may smooth out some of the image-related nuisance variability identified by White and Burge (2025). Integration at a coarser scale is not feasible for stereo, since we have imposed an acuity judgement that requires fine-scale spatial processing and is therefore more vulnerable to the image-related variability, which inevitably differs over a sequence of object views created by the motion. On this argument, stereo may be downweighted by cue combination in a model in which combination of stereo and motion is mandatory (Hillis, Ernst, Banks, & Landy, 2002). Within this model framework, every extra added cue reduces the weight on currently available cues. Alternatively, if stereo can be extracted independently of motion, the variability induced by the sequence of image changes simply makes stereo more noisy.

A third, attentional account is that tracking a moving, rotating object imposes additional processing demands that reduce the resources available for fine binocular analysis, consistent with the longer reaction times we observed. Disentangling these accounts, namely surface non-stationarity, cue competition, and processing load, would require manipulations that vary rotation speed, excursion, and the reliability of the motion-defined shape independently of the disparity signal.

The specificity of the reaction time effect on correct trials is informative in this context. Because incorrect-trial reaction time was flat across disparity in both conditions, the slowing under motion is unlikely to reflect a simple, global increase in response latency; rather, it tracks the detection process itself, suggesting that motion makes the binocular signal genuinely harder to extract rather than merely making observers respond more cautiously. It is worth noting that the threshold effect was both larger and more uniform than the reaction time effect: every observer showed a threshold increase under motion, whereas two of thirteen showed a decrease in reaction time. Accuracy-based threshold estimation appears to be the more sensitive index of the motion cost, with reaction time providing a convergent but noisier measure.

### 4.3 A bridge to neurophysiology

A central motivation for this work was methodological, to establish whether disparity sensitivity can be measured quantitatively with photorealistic objects using a task that could transfer to macaque neurophysiology. The odd-one-out task used here is closely matched to the four-quadrant detection task employed in awake, behaving macaque recordings, in which the animal reports by saccade the location of a target that changes in binocular depth, and in which orderly psychometric and reaction time functions have been obtained alongside simultaneous population recordings from V1 and V4 (Smith & Parker, 2021). The present results show that this design yields orderly psychometric functions and reliable thresholds in humans with photorealistic stimuli, providing both a behavioural benchmark and a validated stimulus set for subsequent recordings.

This is directly relevant to current questions about how mid-level ventral stream areas represent binocular depth in the presence of other cues, and how disparity signals are transformed as they pass through these areas (Parker, 2020; Pasupathy et al., 2020). V4 neurons are disparity selective (Watanabe, Tanaka, Uka, & Fujita, 2002), are tuned for the three-dimensional orientation of contours (Hinkle & Connor, 2002), signal a degree of relative disparity rather than pure, absolute disparity (Umeda, Tanabe, & Fujita, 2007), and carry activity sufficient to support fine disparity discrimination (Shiozaki, Tanabe, Doi, & Fujita, 2012); further downstream, inferotemporal neurons are selective for disparity-defined three-dimensional shape (Janssen, Vogels, & Orban, 1999, 2000). What is not known is how these representations behave when disparity is presented in the context of a fully shaded, recognisable object rather than a random dot stereogram, which is precisely the gap the present stimulus set is designed to address.

The observation that motion substantially alters disparity detection at the behavioural level further suggests a concrete prediction for physiology, the population encoding of disparity in these areas may differ systematically between static and dynamic objects, offering a window into the interactions between the dorsal and ventral streams. Because the correlation structure of population activity, and not only single neuron tuning, limits the information available for depth discrimination (Smith & Parker, 2021), the behavioural cost of motion reported here could arise either from a reduction in single neuron disparity signal or from a change in the correlated variability across the population. The present paradigm, transferred to the macaque, is well placed to distinguish these possibilities.

### 4.4 Limitations and future directions

Several limitations should be acknowledged. A substantial minority of observers, four of sixteen in the static condition and two of thirteen in the motion condition, could not be assigned a threshold within the tested range of disparities. These exclusions did not have a single cause. For two observers (P13, P15), correct-trial reaction time decreased with disparity in the same manner as included observers, and accuracy rose substantially above chance across the tested range (from 25% to 47% correct, and from 20% to 32% correct, respectively), indicating a genuine, if shallower, psychometric function that simply required more disparity than the tested range provided to reach the 62.5% criterion. For the remaining two observers (P14, P16), accuracy remained close to chance across the full range of disparities tested despite an intact correct/incorrect reaction time dissociation, and for both the fitted psychometric function was nearly flat: its slope parameter σ, the standard deviation of the cumulative Gaussian, was 169 and 295 %IPD respectively, larger than the entire stimulus range, so the 62.5% point could not be located. For P14, mean reaction time was also more than twice that of any other observer. These cases are consistent with limited engagement with the task rather than with genuinely poor stereoacuity, whereas the other two are better characterised as the lower tail of a broad population distribution of binocular function (Bosten et al., 2015; Coutant & Westheimer, 1993), which the present disparity range did not extend far enough to capture. Extending the tested range, particularly at the upper end, may allow thresholds to be determined for observers of genuinely lower stereoacuity and would help distinguish these cases from disengagement in future studies, albeit at the expense of making binocular stimuli that do not correspond geometrically to real-world objects. Relating measurements with naturalistic objects to independent clinical measures of stereoacuity would provide a further check. Finally, each object was rendered as if viewed straight ahead, with the cameras converged on its centre, although it was displayed 7.7° from fixation; this departure from exact viewing geometry was the same in both conditions and so does not affect the comparison between them.

The two conditions differed in their tested disparity ranges, reflecting the higher thresholds anticipated for moving objects; while the psychometric fits accommodate this, fully matched ranges would strengthen the comparison. The angular disparities reported here were derived from the rendering and display geometry (Section 2.4). In addition, static sessions always preceded motion sessions, so practice and order are confounded with condition; because practice would be expected to improve performance, this ordering is more likely to have reduced than to have produced the cost of motion, but a counterbalanced design would remove the ambiguity.

Finally, the present design isolates the detection of an added disparity against zero-disparity distractor objects. Future work could parametrically vary the congruence between disparity and the other depth cues, namely shading, texture, and motion, to test directly how these cues are weighted and combined in photorealistic objects, connecting the present detection measures to the cue combination framework (Landy et al., 1995; Johnston et al., 1994; Knill & Saunders, 2003; Hibbard et al., 2025). Care is needed in the design of such experiments, since superficially similar cue-conflict manipulations can fail to distinguish between competing models of integration (Scarfe, 2022). A further natural extension is to exploit the manipulability of the laser-scanned models directly, compressing the depth profile along the line of sight while independently restoring the binocular signal through the rendered camera separation, so that disparity and shape from shading can be traded off against one another quantitatively (Johnston et al., 1993).

### 4.5 Conclusion

Human observers reliably detect binocular disparity carried by photorealistic, laser-scanned objects, with detection thresholds and reaction times that vary in an orderly way with disparity magnitude, and with detection abolished when the binocular disparity of the target is removed. Disparity is detected less well in rotating than in static photorealistic objects, with substantially raised thresholds and slower responses in every observer tested, indicating that object motion imposes a measurable cost on the extraction of binocular depth under naturalistic viewing. Beyond characterising stereopsis in a more ecologically representative setting, these results establish a behavioural foundation and a transferable task design, with depth parameterised in observer-relative units and a decision structure already validated in the macaque, for future neurophysiological investigations of how mid-level visual areas encode binocular depth in realistic visual contexts.

## Notes

### Competing Interest Statement

The authors have declared no competing interest.

### Funding

- Deutsche Forschungsgemeinschaft: 498049560

## References

Baker, D. H., Kaestner, M., & Gouws, A. D. (2016). Measurement of crosstalk in stereoscopic display systems used for vision research. Journal of Vision, 16(15), 14. 10.1167/16.15.14

Bosten, J. M., Goodbourn, P. T., Lawrance-Owen, A. J., Bargary, G., Hogg, R. E., & Mollon, J. D. (2015). A population study of binocular function. Vision Research, 110(Pt A), 34–50. 10.1016/j.visres.2015.02.017

Bradshaw, M. F., & Rogers, B. J. (1996). The interaction of binocular disparity and motion parallax in the computation of depth. Vision Research, 36(21), 3457–3468. 10.1016/0042-6989(96)00072-7

Brainard, D. H. (1997). The Psychophysics Toolbox. Spatial Vision, 10(4), 433–436.

Coutant, B. E., & Westheimer, G. (1993). Population distribution of stereoscopic ability. Ophthalmic and Physiological Optics, 13(1), 3–7.

Cumming, B. G., & DeAngelis, G. C. (2001). The physiology of stereopsis. Annual Review of Neuroscience, 24, 203–238.

Ernst, M. O., & Banks, M. S. (2002). Humans integrate visual and haptic information in a statistically optimal fashion. Nature, 415(6870), 429–433.

Hibbard, P. B., Asher, J. M., & Hornsey, R. L. (2025). The contributions of pictorial, motion, and binocular cues to the perception of depth and distance. Vision Research, 234, 108653.

Hibbard, P. B., Goutcher, R., O’Kane, L. M., & Scarfe, P. (2012). Misperception of aspect ratio in binocularly viewed surfaces. Vision Research, 70, 34–43.

Hibbard, P. B., Hornsey, R. L., & Asher, J. M. (2023). Binocular information improves the reliability and consistency of pictorial relief. Vision, 7(1), 1. 10.3390/vision7010001

Hillis, J. M., Ernst, M. O., Banks, M. S., & Landy, M. S. (2002). Combining sensory information: Mandatory fusion within, but not between, senses. Science, 298(5598), 1627–1630. 10.1126/science.1075396

Hillis, J. M., Watt, S. J., Landy, M. S., & Banks, M. S. (2004). Slant from texture and disparity cues: Optimal cue combination. Journal of Vision, 4(12), 967–992.

Hinkle, D. A., & Connor, C. E. (2002). Three-dimensional orientation tuning in macaque area V4. Nature Neuroscience, 5(7), 665–670.

Hornsey, R. L., Hibbard, P. B., & Scarfe, P. (2015). Ordinal judgements of depth in monocularly and binocularly viewed photographs of complex natural scenes. Proceedings of the International Conference on 3D Imaging (IC3D), 1–6.

Hornsey, R. L., Hibbard, P. B., & Scarfe, P. (2016). Binocular depth judgements on smoothly curved surfaces. PLOS ONE, 11(11), e0165932.

Howard, I. P., & Rogers, B. J. (2002). Seeing in depth (Vols. 1–2). University of Toronto Press.

Jäkel, F., & Wichmann, F. A. (2006). Spatial four-alternative forced-choice method is the preferred psychophysical method for naïve observers. Journal of Vision, 6(11), 13. 10.1167/6.11.13

Janssen, P., Vogels, R., & Orban, G. A. (1999). Macaque inferior temporal neurons are selective for disparity-defined three-dimensional shapes. Proceedings of the National Academy of Sciences, 96(14), 8217–8222.

Janssen, P., Vogels, R., & Orban, G. A. (2000). Three-dimensional shape coding in inferior temporal cortex. Neuron, 27(2), 385–397.

Johnston, E. B., Cumming, B. G., & Landy, M. S. (1994). Integration of stereopsis and motion shape cues. Vision Research, 34(17), 2259–2275.

Johnston, E. B., Cumming, B. G., & Parker, A. J. (1993). Integration of depth modules: Stereopsis and texture. Vision Research, 33(5–6), 813–826.

Julesz, B. (1971). Foundations of cyclopean perception. University of Chicago Press.

Kingdom, F. A. A., & Prins, N. (2016). Psychophysics: A practical introduction (2nd ed.). Academic Press.

Knill, D. C., & Saunders, J. A. (2003). Do humans optimally integrate stereo and texture information for judgments of surface slant? Vision Research, 43(24), 2539–2558.

Landy, M. S., Maloney, L. T., Johnston, E. B., & Young, M. (1995). Measurement and modeling of depth cue combination: In defense of weak fusion. Vision Research, 35(3), 389–412.

McKee, S. P., Levi, D. M., & Bowne, S. F. (1990). The imprecision of stereopsis. Vision Research, 30(11), 1763–1779.

Parker, A. J. (2020). Intermediate level cortical areas and the multiple roles of area V4. Current Opinion in Physiology, 16, 61–67.

Pasupathy, A., Popovkina, D. V., & Kim, T. (2020). Visual functions of primate area V4. Annual Review of Vision Science, 6, 363–385.

Pelli, D. G. (1997). The VideoToolbox software for visual psychophysics: Transforming numbers into movies. Spatial Vision, 10(4), 437–442.

Read, J. C. A., et al. (2020). ASTEROID stereotest v1.0: Lower stereo thresholds using smaller, denser and faster dots. Ophthalmic and Physiological Optics, 40(6), 815–827.

Scarfe, P. (2022). Experimentally disambiguating models of sensory cue integration. Journal of Vision, 22(1), 5. 10.1167/jov.22.1.5

Scarfe, P., & Glennerster, A. (2014). Humans use predictive kinematic models to calibrate visual cues to three-dimensional surface slant. Journal of Neuroscience, 34(31), 10394–10401.

Scarfe, P., & Glennerster, A. (2015). Using high-fidelity virtual reality to study perception in freely moving observers. Journal of Vision, 15(9), 3. 10.1167/15.9.3

Scarfe, P., & Glennerster, A. (2019). The science behind virtual reality displays. Annual Review of Vision Science, 5, 529–547.

Scarfe, P., & Hibbard, P. B. (2006). Disparity-defined objects moving in depth do not elicit three-dimensional shape constancy. Vision Research, 46(10), 1599–1610.

Scarfe, P., & Hibbard, P. B. (2011). Statistically optimal integration of biased sensory estimates. Journal of Vision, 11(7), 12. 10.1167/11.7.12

Scarfe, P., & Hibbard, P. B. (2013). Reverse correlation reveals how observers sample visual information when estimating three-dimensional shape. Vision Research, 86, 115–127.

Serrano-Pedraza, I., Vancleef, K., & Read, J. C. A. (2016). Avoiding monocular artifacts in clinical stereotests presented on column-interleaved digital stereoscopic displays. Journal of Vision, 16(14), 13. 10.1167/16.14.13

Shiozaki, H. M., Tanabe, S., Doi, T., & Fujita, I. (2012). Neural activity in cortical area V4 underlies fine disparity discrimination. The Journal of Neuroscience, 32(11), 3830–3841.

Smith, J. E. T., & Parker, A. J. (2021). Correlated structure of neuronal firing in macaque visual cortex limits information for binocular depth discrimination. Journal of Neurophysiology, 126(1), 275–303.

Tittle, J. S., & Braunstein, M. L. (1993). Recovery of 3 D shape from binocular disparity and structure-from-motion. Perception & Psychophysics, 54(2), 157–169.

Tittle, J. S., & Perotti, V. J. (1997). The perception of shape and curvedness from binocular stereopsis and structure-from-motion. Perception & Psychophysics, 59(8), 1167–1179.

Umeda, K., Tanabe, S., & Fujita, I. (2007). Representation of stereoscopic depth based on relative disparity in macaque area V4. Journal of Neurophysiology, 98(1), 241–252.

Vancleef, K., Read, J. C. A., Herbert, W., Goodship, N., Woodhouse, M., & Serrano-Pedraza, I. (2018). Two choices good, four choices better: For measuring stereoacuity in children, a four-alternative forced choice paradigm is more efficient than two. PLOS ONE, 13(7), e0201366.

Walraven, J. (n.d.). TNO test for stereoscopic vision. Lameris Ootech.

Watanabe, M., Tanaka, H., Uka, T., & Fujita, I. (2002). Disparity selective neurons in area V4 of macaque monkeys. Journal of Neurophysiology, 87(4), 1960–1973.

White, D. N., & Burge, J. (2025). How distinct sources of nuisance variability in natural images and scenes limit human stereopsis. PLOS Computational Biology, 21(4), e1012945. 10.1371/journal.pcbi.1012945

Wichmann, F. A., & Hill, N. J. (2001a). The psychometric function: I. Fitting, sampling, and goodness of fit. Perception & Psychophysics, 63(8), 1293–1313.

Wichmann, F. A., & Hill, N. J. (2001b). The psychometric function: II. Bootstrap-based confidence intervals and sampling. Perception & Psychophysics, 63(8), 1314–1329.

World Medical Association. (2025). World Medical Association Declaration of Helsinki: Ethical principles for medical research involving human participants. JAMA, 333(1), 71–74. 10.1001/jama.2024.21972

