## Supplemental Material for "Detecting Binocular Disparity in Photorealistic Objects"

#### S1. Per-observer summary

Because the number of observers contributing to each analysis differs, this table records every observer's data and which analyses they enter. Sixteen observers completed the static condition; thirteen of these also completed the motion condition. Thresholds could not be estimated within the tested range for four observers in the static condition and two in the motion condition, under the pre-specified rule described in the Methods.

| Obs | Static %IPD | arcmin | Static status | Motion %IPD | arcmin | Motion status | Static RT (ms) | Motion RT (ms) |
|---|---|---|---|---|---|---|---|---|
| P1 | 66.83 | 1.06 | included | 144.77 | — | excluded | 2112.4 | 1498.5 |
| P2 | 49.85 | 0.79 | included | 99.93 | 1.58 | included | 1792.8 | 2221.5 |
| P3 | 34.40 | 0.54 | included | 84.68 | 1.34 | included | 1842.7 | 3180.9 |
| P4 | 50.39 | 0.80 | included | 75.53 | 1.19 | included | 2635.2 | 2991.3 |
| P5 | 54.39 | 0.86 | included | 89.60 | 1.42 | included | 1970.8 | 2177.8 |
| P6 | 55.95 | 0.87 | included | 89.25 | 1.39 | included | 2765.1 | 3782.8 |
| P7 | 42.15 | 0.66 | included | 85.30 | 1.33 | included | 3046.4 | 3319.7 |
| P8 | 89.49 | 1.30 | included | 131.04 | 1.91 | included | 2680.9 | 3027.1 |
| P9 | 73.98 | 1.15 | included | 106.75 | 1.66 | included | 3039.8 | 3687.8 |
| P10 | 44.15 | 0.69 | included | 78.66 | 1.22 | included | 3235.1 | 4011.9 |
| P11 | 49.59 | 0.77 | included | 97.50 | 1.52 | included | 2930.1 | 3299.4 |
| P12 | 79.79 | 1.24 | included | 113.57 | 1.77 | included | 2867.7 | 2696.4 |
| P13 | 128.62 | — | excluded | 200.87 | — | excluded | 3124.2 | 3738.2 |
| P14 | 1470.72 | — | excluded | — | — | not run | 7459.8 | — |
| P15 | 188.88 | — | excluded | — | — | not run | 3786.4 | — |
| P16 | 364.88 | — | excluded | — | — | not run | 2948.4 | — |

*Note.* Thresholds are maximum-likelihood estimates of the effective %IPD yielding 62.5% correct, with the equivalent maximum within-object disparity in arcmin of visual angle, converted at each observer's own rendered interpupillary distance (S2). Angular values are given only where the threshold lies within the tested range; for excluded observers the %IPD figure is an extrapolation and no angular equivalent is meaningful. Reaction times are means over correct trials, pooled across disparity levels and excluding control trials. For P6, P7 and P9–P13, whose motion stimuli were rendered at 63 mm rather than at the measured interpupillary distance, motion thresholds are expressed as a percentage of the measured interpupillary distance (nominal %IPD × 63 / measured interpupillary distance), as in the main text. Analyses drawing on this table: static thresholds $n$ = 12; motion thresholds $n$ = 11; paired threshold comparison $n$ = 11 (P2–P12); paired reaction-time comparison $n$ = 13 (P1–P13); static reaction-time ANOVA $n$ = 16; motion reaction-time ANOVA $n$ = 13.

#### S2. Disparity calibration

Depth magnitude was specified as a percentage of each observer's interpupillary distance, so the same nominal level corresponds to a different absolute disparity in different observers and, because the four objects differ in depth profile, to slightly different disparities across objects. This section gives the conversion in full.

#### S2.1 Conversion constants

Scene disparity was obtained from the rendering geometry as $\delta = b_{eff} \cdot k$, where $b_{eff}$ is the effective camera baseline and $k = 1/Z_S - 1/Z_C$ with $Z_C$ the convergence distance and $Z_S$ the distance to the nearest surface point of the object. Because the images depend only on the ratios between these lengths, all scene lengths are expressed in units of the observer's interpupillary distance (IPD; main text, Section 2.4). The baseline in units of IPD equals the rendering level divided by 100 and multiplied by the gain $g$ (for example 0.4 IPD at 40 %IPD with $g = 1$), with the two cameras displaced symmetrically about the central viewpoint, and for a 63 mm interpupillary distance $Z_C$ is 96.17, 97.67, 87.18 and 84.10 IPD and $Z_S$ is 86.45, 88.56, 78.50 and 75.97 IPD for the apple, lemon, nectarine and garlic. Converting to pixels uses the focal length $f_{PX}$ = 2666.7 px. Retinal disparity then follows from the display geometry: one rendered pixel is drawn at 0.59 of native size on a projected image 424 mm wide at 1920 px, viewed from 900 mm, giving 0.1303 mm on screen and 0.4977 arcmin at the eye. For comparison, the angle subtended at the rendering camera is 1.2892 arcmin per rendered pixel, 2.59 times larger; the camera figure is not the retinal disparity and is not used here.

#### S2.2 Angular disparity per unit %IPD, by interpupillary distance and object

| IPD (mm) | apple | garlic | lemon | nectarine | Mean | CV |
|---|---|---|---|---|---|---|
| 58 | 0.014280 | 0.015542 | 0.012867 | 0.015510 | 0.014550 | 0.087 |
| 59 | 0.014527 | 0.015810 | 0.013089 | 0.015777 | 0.014801 | 0.087 |
| 60 | 0.014773 | 0.016078 | 0.013311 | 0.016044 | 0.015052 | 0.087 |
| 61 | 0.015019 | 0.016346 | 0.013533 | 0.016312 | 0.015302 | 0.087 |
| 62 | 0.015265 | 0.016614 | 0.013755 | 0.016579 | 0.015553 | 0.087 |
| 63 | 0.015511 | 0.016882 | 0.013976 | 0.016847 | 0.015804 | 0.087 |

*Note.* Values are arcmin of retinal disparity per unit of effective %IPD. The scene coefficient $k = 1/Z_S - 1/Z_C$ is a property of the object: for a 63 mm interpupillary distance it is 0.001169 (apple), 0.001272 (garlic), 0.001053 (lemon) and 0.001269 (nectarine) rad per IPD of baseline, and it scales with IPD / 63 mm because the rendered scene was the same for every observer. Disparity scales linearly with interpupillary distance, so an observer rendered at 63 mm receives 8.6% more absolute disparity than one at 58 mm for the same nominal level. Between objects the coefficient of variation is 0.087 at every interpupillary distance, so a nominal level corresponds to a range of angular disparities spanning roughly ±9% about the mean rather than to a single value. Angular thresholds in the main text are means across the four objects, computed at each observer's own rendered interpupillary distance.

#### S2.3 Angular disparity at each tested level

| %IPD | 58 mm | 59 mm | 60 mm | 61 mm | 62 mm | 63 mm |
|---|---|---|---|---|---|---|
| 0 | 0.00 | 0.00 | 0.00 | 0.00 | 0.00 | 0.00 |
| 20 | 0.29 | 0.30 | 0.30 | 0.31 | 0.31 | 0.32 |
| 40 | 0.58 | 0.59 | 0.60 | 0.61 | 0.62 | 0.63 |
| 60 | 0.87 | 0.89 | 0.90 | 0.92 | 0.93 | 0.95 |
| 80 | 1.16 | 1.18 | 1.20 | 1.22 | 1.24 | 1.26 |
| 100 | 1.46 | 1.48 | 1.51 | 1.53 | 1.56 | 1.58 |
| 125 | 1.82 | 1.85 | 1.88 | 1.91 | 1.94 | 1.98 |

*Note.* Maximum within-object disparity in arcmin of visual angle, averaged across the four objects. The 125 %IPD level was presented in the motion condition only.

#### S3. Excluded observers

Four observers in the static condition could not be assigned a threshold within the tested range. To establish whether these exclusions reflected disengagement from the task or genuinely low sensitivity, two properties were examined that do not depend on the psychometric fit converging: whether correct-trial reaction time decreased with disparity, as it did for every included observer, and whether accuracy rose above chance across the tested range.

##### S3.1 Correct-trial reaction-time slope

| Observer | Status | Slope (ms per %IPD) |
|---|---|---|
| P13 | excluded | −10.96 |
| P14 | excluded | −47.46 |
| P15 | excluded | −6.31 |
| P16 | excluded | −21.99 |
| P1 | included | −6.86 |
| P2 | included | −5.43 |
| P3 | included | −7.73 |
| P6 | included | −18.06 |
| P7 | included | −42.04 |
| P9 | included | −30.77 |

*Note.* All four excluded observers show the negative slope characteristic of engaged responding, within the range spanned by included observers. Reaction time alone therefore does not distinguish them.

##### S3.2 Accuracy by disparity level

| Observer | Status | 0 | 20 | 40 | 60 | 80 | 100 |
|---|---|---|---|---|---|---|---|
| P13 | excluded | 25.0 | 23.3 | 26.7 | 38.3 | 36.7 | 46.7 |
| P14 | excluded | 20.0 | 23.3 | 26.7 | 23.3 | 25.0 | 26.7 |
| P15 | excluded | 20.0 | 18.3 | 28.3 | 35.0 | 28.3 | 31.7 |
| P16 | excluded | 30.0 | 36.7 | 35.0 | 40.0 | 40.0 | 35.0 |
| P1 | included | 30.0 | 26.7 | 46.7 | 61.7 | 71.7 | 81.7 |
| P2 | included | 28.3 | 33.3 | 56.7 | 78.3 | 86.7 | 85.0 |
| P3 | included | 23.3 | 46.7 | 78.3 | 80.0 | 96.7 | 96.7 |
| P6 | included | 16.7 | 31.7 | 50.0 | 80.0 | 81.7 | 83.3 |
| P7 | included | 18.3 | 40.0 | 68.3 | 86.7 | 81.7 | 96.7 |
| P9 | included | 25.0 | 30.0 | 40.0 | 51.7 | 61.7 | 85.0 |

*Note.* Percent correct; chance is 25%. Accuracy separates the excluded observers into two groups. P13 and P15 rose substantially above chance (25% to 47%, and 20% to 32%), indicating a real psychometric function whose criterion simply lay beyond the disparities presented. P14 and P16 remained near chance at every level despite the full range being tested, and both had fitted slope parameters exceeding that range (169.45 and 294.97 %IPD). These two are consistent with limited engagement rather than with low but genuine stereoacuity.

### S4. Sensitivity analyses

#### S4.1 Thresholds with and without the exclusion rule

| Condition | Rule | n | Mean %IPD | SD | Median | Range |
|---|---|---|---|---|---|---|
| Static | Applied | 12 | 57.58 | 16.54 | 52.39 | 34.40–89.49 |
| Static | Not applied | 16 | 177.75 | 354.55 | 61.39 | 34.40–1470.72 |
| Motion | Applied | 11 | 95.62 | 16.48 | 89.60 | 75.53–131.04 |
| Motion | Not applied | 13 | 107.50 | 34.61 | 97.50 | 75.53–200.87 |

*Note.* Without the exclusion rule the static mean is inflated more than threefold and the standard deviation exceeds the mean by a factor of two, driven by fits whose criterion was never bracketed by the data. The median is comparatively stable across rules (52.39 against 61.39), which is itself diagnostic that the excluded values are extrapolations rather than the upper tail of a continuous distribution.

#### S4.2 Reaction-time ANOVA with and without near-chance observers

| Analysis | n | F | df | p | partial $\eta^2$ | GG ε | GG p |
|---|---|---|---|---|---|---|---|
| Static, all | 16 | 12.17 | 5, 75 | < 0.001 | 0.448 | 0.409 | < 0.001 |
| Static, excl. P14/P16 | 14 | 12.83 | 5, 65 | < 0.001 | 0.497 | 0.268 | 0.001 |
| Motion, all | 13 | 9.15 | 6, 72 | < 0.001 | 0.433 | 0.348 | < 0.001 |

*Note.* Correct-trial reaction time. Removing the two near-chance observers slightly strengthens the effect rather than weakening it, so the reported result does not depend on their inclusion. Sphericity is strongly violated throughout, reflecting greater variance in reaction time at low disparity; Greenhouse-Geisser corrected values are reported alongside.

### S5. Individual psychometric functions

Fitted cumulative Gaussian functions for every observer in each condition, including those whose threshold could not be estimated within the tested range. Symbols are the observed proportion correct at each effective %IPD, curves the maximum-likelihood fits, shaded bands the central 95% of fitted functions across 400 parametric-bootstrap samples, and vertical dashed lines the fitted threshold (labelled in %IPD) where it lies within the range presented. Colours and symbols for P1–P12 match Figures 4, 5 and 8 of the main text. The horizontal dashed line marks chance (25%) and the dotted line the 62.5% criterion.

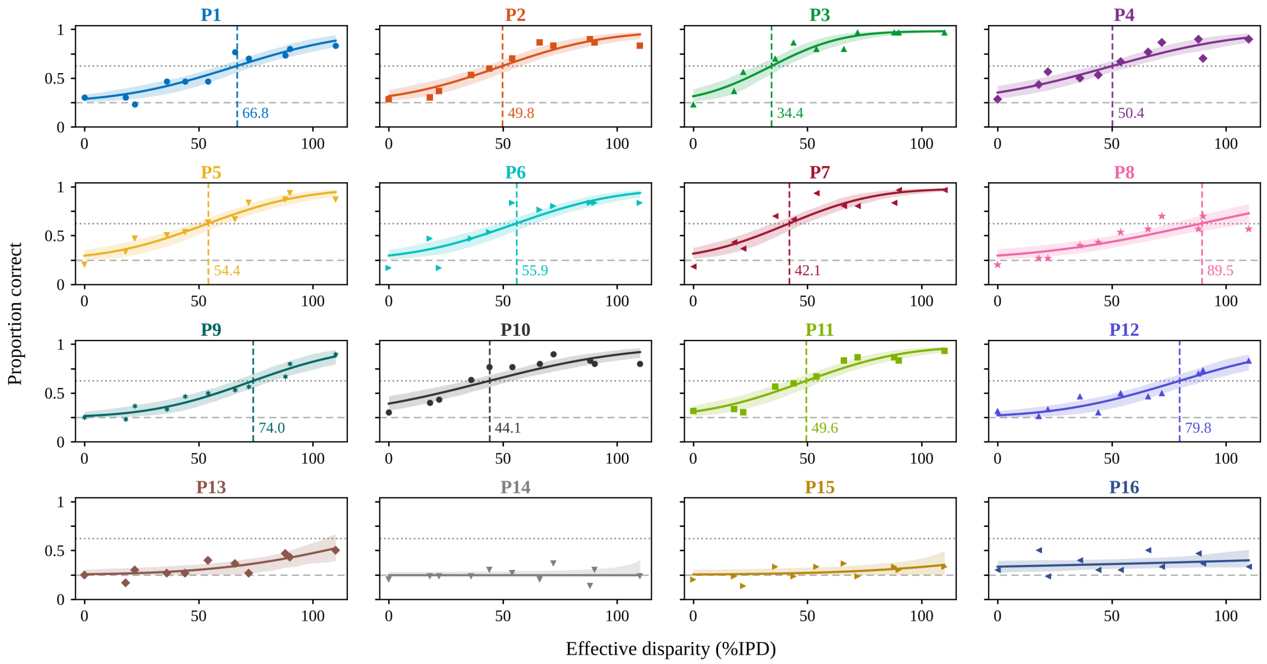


**Figure S1.** *Static condition, all sixteen observers.* Twelve observers (P1–P12) yielded a threshold within the tested range, shown as a vertical dashed line. For the remaining four no threshold line is drawn, because the fitted criterion lay beyond the disparities presented. The shapes of these four functions differ in a way that matters for interpretation: P13 and P15 rise steadily above chance across the range, indicating a real but shallow psychometric function, whereas P14 is flat at chance throughout and P16 varies around 0.35 without systematic dependence on disparity. This is the visual counterpart of the accuracy data in S3.2.

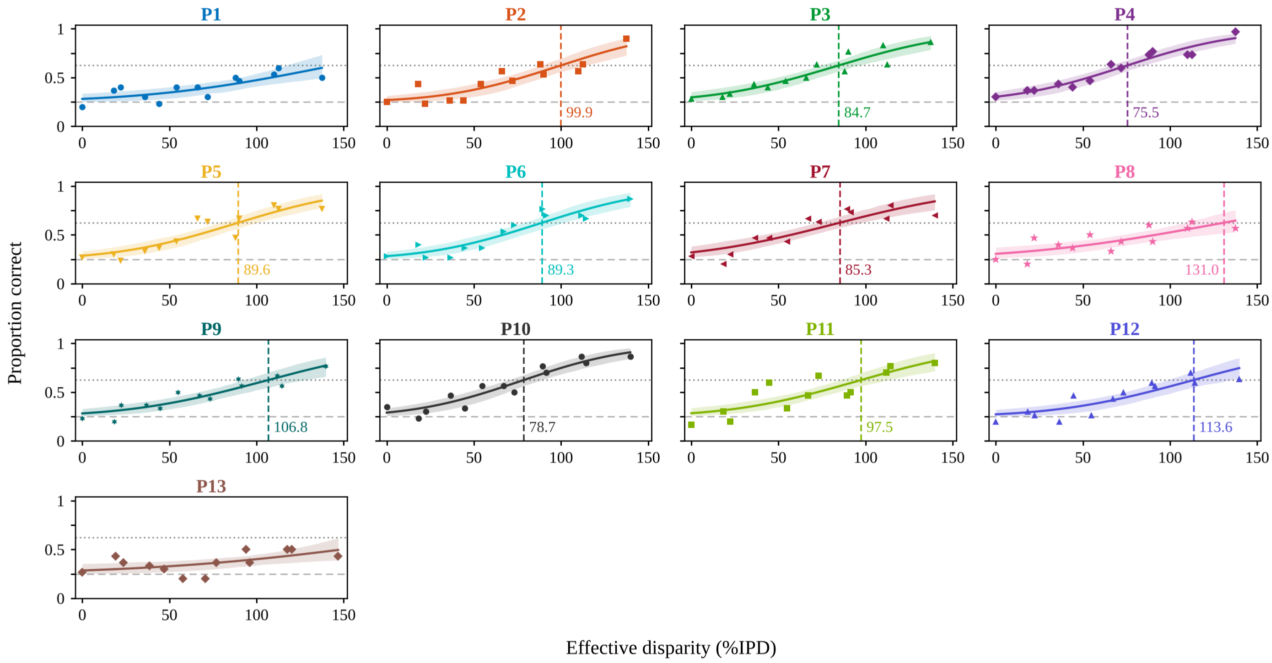


**Figure S2.** *Motion condition, all thirteen observers.* Eleven observers yielded a threshold within the tested range. P1 and P13 did not: P1 rises to approximately 0.6 correct at the highest disparity without crossing the criterion, and P13 rises only to about 0.45 correct at the highest disparities. Note that the disparity axis extends to 137.5–146.8 %IPD here, against 110 %IPD in the static condition, reflecting the additional 125 %IPD level included in this condition.

#### S6. Reaction-time aggregation

Reaction times were aggregated by averaging within observer before averaging across observers, matching the repeated-measures analysis reported in the main text. An alternative is to pool all trials from all observers, but this weights observers by their number of correct trials, which is not constant across disparity levels: at 0 %IPD every observer is at chance and contributes a similar number of correct trials, whereas at high disparity the more sensitive observers contribute far more. Because more sensitive observers also respond faster, trial pooling progressively over-weights the fastest observers as disparity increases, steepening the apparent decline.

##### S6.1 Static condition

| %IPD | Participant-mean (ms) | Trial-pooled (ms) | Difference | Correct trials per observer |
|---|---|---|---|---|
| 0 | 4111.5 | 4033.1 | 78.4 | 14.9 |
| 20 | 3944.9 | 3800.7 | 144.2 | 19.9 |
| 40 | 3386.6 | 3175.0 | 211.6 | 28.8 |
| 60 | 2874.9 | 2650.3 | 224.5 | 36.3 |
| 80 | 2581.9 | 2304.2 | 277.8 | 40.5 |
| 100 | 2461.2 | 2202.9 | 258.2 | 43.1 |

*Note.* $n = 16$. The discrepancy grows from 78 ms at 0 %IPD to 278 ms at 80 %IPD, tracking the rise in correct trials per observer in the final column. The participant-mean values are those plotted in Figure 7A and tested in the reported ANOVA.

##### S6.2 Motion condition

| %IPD | Participant-mean (ms) | Trial-pooled (ms) | Difference | Correct trials per observer |
|---|---|---|---|---|
| 0 | 4086.4 | 4251.6 | −165.2 | 15.4 |
| 20 | 3625.0 | 3586.2 | 38.9 | 18.8 |
| 40 | 3326.4 | 3410.4 | −84.0 | 22.3 |
| 60 | 3103.7 | 3101.9 | 1.9 | 27.2 |
| 80 | 2964.9 | 3019.0 | −54.1 | 34.1 |
| 100 | 2707.1 | 2727.9 | −20.8 | 38.3 |
| 125 | 2574.2 | 2587.6 | −13.4 | 42.3 |

*Note.* $n = 13$. Unlike the static condition, the two aggregation methods agree closely here, with differences that are small and unsystematic in sign. The systematic bias in the static condition therefore reflects the particular relationship between sensitivity and speed among those observers rather than a general property of the paradigm.

#### S7. Monocular control

A control tested whether the target object could be located by anything other than its binocular disparity. On control trials the target was shown with its left-eye image in both eyes, so that it carried no binocular disparity; the three distractor objects were, as on every trial, the left-eye image of the stereo pair shown to both eyes. Control trials (40 per observer) were interleaved with the main trials of static sessions and were not assigned a %IPD level. M2, M3 and M4 are P15, P12 and P16; M1 is a further control session run separately from the main experiment.

| Observer | Correct | Trials | % correct | 95% CI | p vs 0.25 |
|---|---|---|---|---|---|
| M1 | 9 | 40 | 22.5 | [10.8, 38.5] | 0.856 |
| M2 | 12 | 40 | 30.0 | [16.6, 46.5] | 0.467 |
| M3 | 10 | 40 | 25.0 | [12.7, 41.2] | 1.000 |
| M4 | 11 | 40 | 27.5 | [14.6, 43.9] | 0.716 |
| Pooled | 42 | 160 | 26.2 | [19.6, 33.8] | 0.715 |

*Note.* Confidence intervals are exact; $p$ values are from two-sided exact binomial tests against the 25% chance baseline. No observer differed from chance individually, and neither did the pooled estimate. The upper confidence bound on pooled performance is 33.8%, so accuracy above that level can be excluded; the null result reflects adequate sensitivity rather than an uninformative test.

### S8. Comparison with random-dot cylinders

Four observers (P4, P5, P9 and P12) also completed a spatial four-alternative odd-one-out task with random-dot stereograms of rotating cylinders, presented at the same four positions as the fruit. The task was intended to present disparities matched to those of the fruit stimuli, so that the same observers could be compared on a conventional stereo stimulus. Each cylinder comprised 350 dots (3 px) rotating at 65°/s; on target trials the whole cylinder was displaced in depth by a disparity step, rather than carrying added depth structure within the object as the fruit did. The comparison is reported here rather than in the main text because the two tasks differed in several respects, described below.

The disparities presented differed from their nominal labels for two reasons: the disparity values were read from a table that lagged the intended rendering levels by one level, and the presentation script assumed a screen width of 530 mm rather than the actual 424 mm. Cylinder thresholds were therefore fitted against the disparity actually presented, computed from the pixel disparity on each trial at 0.8434 arcmin per screen pixel, using the same maximum-likelihood cumulative Gaussian fit as in the main analysis (guess rate 0.25, lapse rate 0.02, 62.5% correct criterion). All cylinder disparities were computed for a 62 mm interpupillary distance.

| Observer | Cylinder (arcmin) | Fruit, static (arcmin) | Fruit, rotating (arcmin) |
|---|---|---|---|
| P4 | 0.66 | 0.80 | 1.19 |
| P5 | 1.94 | 0.86 | 1.42 |
| P9 | 3.38 | 1.15 | 1.66 |
| P12 | ≈ 4.0* | 1.24 | 1.77 |

*Note.* Thresholds are in arcmin at 62.5% correct: for the fruit, the maximum within-object disparity; for the cylinders, the disparity of the depth step. Fruit values are from Table 1 of the main text. *The cylinder threshold for P12 lay at the upper edge of the tested range and is approximate. Observer rank order was the same for all three measures (P4 < P5 < P9 < P12). Because the cylinder task differed from the fruit task in stimulus class, in the form of the disparity signal (a step in depth of the whole object rather than depth within it), in dot density and in rotation speed, absolute values should not be compared across tasks.

### S9. Per-observer model comparison and goodness of fit

For each observer who completed both conditions, the static and motion data were fitted jointly with two nested models: a fuller model with separate thresholds for the two conditions, and a lesser model in which both conditions shared one threshold. Both models allowed each condition its own slope. The likelihood ratio between the models was compared with its distribution under the lesser model, obtained by refitting both models to 500 simulated data sets. Goodness of fit of each single-condition fit was assessed from its deviance, compared with the deviances of 1,000 data sets simulated from the fitted function.

| Obs | Static %IPD | Motion %IPD | Likelihood ratio | p (Monte Carlo) | Static fit p | Motion fit p |
|---|---|---|---|---|---|---|
| P1* | 66.8 | 144.8 | 37.9 | ≤ 0.002 | 0.629 | 0.547 |
| P2 | 49.8 | 99.9 | 51.3 | ≤ 0.002 | 0.376 | 0.391 |
| P3 | 34.4 | 84.7 | 51.9 | ≤ 0.002 | 0.272 | 0.776 |
| P4 | 50.4 | 75.5 | 12.9 | ≤ 0.002 | 0.334 | 0.892 |
| P5 | 54.4 | 89.6 | 27.0 | ≤ 0.002 | 0.499 | 0.409 |
| P6 | 55.9 | 89.3 | 24.1 | ≤ 0.002 | 0.004 | 0.621 |
| P7 | 42.1 | 85.3 | 34.2 | ≤ 0.002 | 0.012 | 0.151 |
| P8 | 89.5 | 131.0 | 6.6 | 0.010 | 0.170 | 0.392 |
| P9 | 74.0 | 106.8 | 15.9 | ≤ 0.002 | 0.938 | 0.819 |
| P10 | 44.1 | 78.7 | 23.4 | ≤ 0.002 | 0.081 | 0.684 |
| P11 | 49.6 | 97.5 | 46.4 | ≤ 0.002 | 0.789 | 0.021 |
| P12 | 79.8 | 113.6 | 11.5 | ≤ 0.002 | 0.826 | 0.362 |
| P13* | 128.6 | 200.9 | 2.3 | 0.122 | 0.862 | 0.359 |

*Note.* Thresholds are maximum-likelihood estimates (%IPD). With 500 simulations the smallest attainable Monte Carlo p is 0.002. All 11 observers in the paired comparison (P2–P12) show a significantly higher threshold for motion than for static objects; summed over these observers, the likelihood ratio is 305.0 ($\chi^2(11)$, $p < 0.001$). *P1 and P13 are not in the paired comparison because at least one of their thresholds lay outside the tested range; P1 shows a significant difference in the same direction and P13 a non-significant one. Static fit p and motion fit p are goodness-of-fit p values (proportion of simulated deviances at least as large as the observed deviance); values below 0.05 (P6 and P7 static, P11 motion) indicate more scatter between levels than the binomial model predicts. Fits for P14–P16, who completed the static condition only, also gave acceptable deviances ($p = 0.785$, 0.645 and 0.352).
